# m6A RNA modification of m*Htt* intron 1 regulates the generation of *Htt1a* in Huntington’s Disease

**DOI:** 10.1101/2023.11.10.566530

**Authors:** Anika Pupak, Irene Rodríguez Navarro, Kirupa Sathasivam, Amelie Essmann, Ankita Singh, Daniel del Toro, Silvia Ginés, Gillian P. Bates, Ulf Andersson Vang Ørom, Eulalia Marti, Verónica Brito

## Abstract

Huntington’s disease (HD) is a dominantly inherited neurodegenerative disorder caused by an expanded, somatically unstable CAG repeat in the first exon of the huntingtin gene (*HTT)*. In the presence of an expanded CAG repeat, huntingtin mRNA undergoes an aberrant processing that generates *HTT1a* transcripts with exon 1 and intron 1 sequences, which encodes the aggregation-prone and pathogenic HTTexon 1 protein. The regulatory mechanisms that contribute to the production of *HTT1a* are not fully understood. In a previous transcriptome-wide m6A landscape study performed in *Hdh^+/Q111^* knock-in mice, we have found that the proximal region of intron 1 to exon1-intron 1 splice site in *Htt* RNA is highly modified by m6A. Several pieces of evidence have demonstrated that m6A is involved in RNA processing and splicing. Therefore, in this study we set out to explore the impact of m6A RNA modifications in the generation of *Htt1a*. We show in the striatum of *Hdh^+/Q111^* mice that m6A is enriched in intronic sequences 5’ to the cryptic poly (A) sites (IpA1 and IpA2) at 680 and 1145 bp into intron 1 as well as in *Htt1a* polyadenylated mRNA. We also verified the presence of specific m6A-modified sites near the 5’ exon1-intron1 splice donor site. Intronic *HTT* m6A methylation was recapitulated in human samples showing a significantly increased methylation ratio in HD putamen *post-mortem* samples and in HD fibroblast cell lines from pre-symptomatic and symptomatic patients. In order to test the hypothesis that the m6A modification is involved in mutant *Htt* RNA processing, we performed a pharmacological inhibition of METTL3 and a targeted demethylation of *Htt* intron 1 in HD cells using a dCas13-ALKBH5 system. We found that *Htt1a* transcript levels in HD cells are regulated by METTL3 and by methylation status in *Htt* intron 1. Site-specific manipulation with an RNA editing system resulted in decreased expression levels of *Htt1a*, which was accompanied by a reduction in DNA damage, a major hallmark in HD. Finally, we propose that m6A methylation in intron 1 is likely dependent on the expanded CAG repeats. These findings provide insight into the role of m6A in the generation of the aberrantly spliced mutant *Htt* transcripts with important implications for therapeutic strategies.

## INTRODUCTION

Huntington’s disease (HD) is considered the most common monogenetic neurodegenerative disorder showing dominant inheritance. This disorder is characterized by a triad of motor, cognitive and psychiatric symptoms which largely affect patients’ quality of life ^1^, eventually leading to their death 15-20 years after the diagnosis ^2^. HD is caused by an unstable CAG repeat expansion in exon 1 of the gene that encodes for huntingtin (*HTT*), resulting in an abnormally long polyglutamine tract in the huntingtin protein (HTT) which causes protein misfolding and consequent neurotoxicity ^3^. In the context of expanded CAGs, it has been described that besides full length (FL) *HTT* mRNA isoforms, two small transcripts containing exon 1 and a 5’ region of intron 1 sequences (*HTT1a*) can be generated by incomplete splicing due to aberrant polyadenylation at cryptic polyA sites within intron 1 followed by premature termination of transcription ^4,5^. *HTT1a* not only encodes the aggregation-prone HTTexon1 protein that is known to be highly pathogenic ^6^ but it can also form mRNA nuclear clusters which are resistant to treatment with *HTT* antisense oligonucleotides (ASOs) ^7,8^. As the alternative processing of *HTT* mRNA, and consequently the levels of *HTT1a* and HTTexon1, increase with increasing CAG repeat length ^9^ it has been suggested that *HTT1a* and HTTexon1 may be the pathogenic contributor of somatic CAG repeat expansion in HD ^10^. Several regulatory mechanisms influencing the amount of *HTT1a* production have been proposed, such as RNA polymerase II (PolII) transcription speed and binding of splicing factors to the expanded CAG repeats or intron 1 of *HTT* mRNA ^4,11, 12^. However, the impact of RNA modifications on the transcriptional and post-transcriptional regulation of *HTT* has so far not been explored.

N(6)-methyladenosine (m6A) is the most abundant internal modification in eukaryotic mRNA ^13, 14^, being present in 0.2-0.6% of all adenosines in mammalian mRNA ^15^. The discovery of the m6A methyltransferase complex (METTL3, METTL14 and WTAP) and the demethylase proteins (FTO and ALKBH5) (writer and erasers, respectively) showed that this modification exhibits a dynamic pattern, strengthening its regulatory role in gene expression control ^16^. Indeed, m6A has been shown to influence several steps of RNA metabolism as transcription of the nascent RNA, including alternative splicing and polyadenylation, nuclear export, translation and finally degradation ^17–21^, processes that are mediated by the m6A-reader proteins. Notably, we have recently demonstrated in the HD hippocampus that alterations of m6A modifications occur in the mRNA along progression and that these alterations are involved in the cognitive disturbances of *Hdh^+/Q111^* mice ^22^. One crucial finding of this work was a pronounced differential methylation in the proximal region of intron 1 of huntingtin transcripts. Moreover, m6A deposited in introns slow down kinetic the mRNA processing ^23^ and can impact termination of PolII ^24, 25^. Altogether, emphasize the need of a deeper characterization of the effects of m6A on the processing of *HTT* RNA. Therefore, to get more insight into the potential role of m6A in HD pathology, we studied the *Htt* m6A RNA modification in the striatum of HD samples and explored its contribution to *Htt1a* generation. Our findings indicate that the m6A methylation is present in intron 1 of mutant huntingtin (*mHtt*) in the striatum of *Hdh^+/Q111^* mice, further being maintained upon maturation of the RNA. Moreover, we confirmed methylation in specific m6A sites in intron 1 of different HD *in vitro* cell models and we identified a GGACA motif present in human intron 1 to be differentially methylated in human HD fibroblasts and *post-mortem* samples, highlighting its pathological relevance. Notably, pharmacological inhibition of METTL3 or targeted demethylation of *Htt* intron 1 specifically decreases transcript levels of *Htt1a* in HD cells. We further demonstrate that m6A methylation in intron 1 is likely dependent on CAG expansions. Collectively, our findings support the involvement of m6A in the generation of aberrantly spliced *Htt1a*, which could have important implications for gene therapy strategies designed to specifically lower *mHTT* in HD patients.

## METHODS

### *Post-mortem* brain tissue

Human *post-mortem* samples derived from putamen (11 controls and 13 HD patients) were obtained from the Neurological Tissue Banc (Biobanc-Hospital Clínic-Institut d’Investigacions Biomèdiques August Pi I Sunyer (IDIBAPS) according to the guidelines and approval of Barcelona’s Clinical research Ethical Committee (Hospital Clínic). All the ethical guidelines contained within the latest Declaration of Helsinki were taken into consideration, and informed consent was obtained from all subjects under study. Clinical details of controls and HD patients are summarized in Supplementary Table 1.

### Human skin fibroblasts

Human skin fibroblasts were obtained from controls and HD patients at different clinical stages in collaboration with Dr Jaime Kulisevsky and Dr Jesús Pérez from the Hospital de la Santa Creu I San Pau de Barcelona (7 controls, 7 pre-symptomatic HD patients and 12 symptomatic HD patients). All procedures were approved by the Ethics Committees of the Hospital de la Santa Creu I San Pau de Barcelona and the Universitat de Barcelona and informed written consent was obtained for all subjects. Clinical data of the subjects is summarized in Supplementary table 2. Cells derived from sterile, non-necrotic skin biopsies were grown at 37 °C and 5% CO_2_ in Dulbecco’s modified Eagle’s medium (DMEM) with 25 mM glucose (Gibco, ref. 41966-029) supplemented with 10% fetal bovine serum (FBS), 1% penicillin-streptomycin and 1% amphotericin B.

### Animals

Heterozygous *Hdh^+/Q111^* knock-in mice ^26^ were used as an HD mouse model. Those mice were maintained on a C57BL/6J (Charles River) genetic background and present a targeted insertion of a 109 CAG repeats in the murine huntingtin gene that extends the resulting polyglutamine segment to 111 residues. Male *Hdh^Q7/Q7^*WT mice were crossed with female heterozygous *Hdh^+/Q111^* mice to obtain age matched WT and *Hdh^+/Q111^* littermates. Only males from each genotype were analyzed. Mice were housed with access to food and water *ad libitum* in a colony room kept at 19-22°C and 40-60% humidity, under a 12:12 h light/dark cycle. Animals were sacrificed at 2 and 8 months of age through cervical dislocation, and brains were rapidly harvested and dissected, frozen in dry ice and stored at −80°C.

### Immortalized cell culture and pharmacological inhibition

Conditionally immortalized murine homozygous wild-type ST*Hdh^Q7/Q7^* and mutant ST*Hdh^Q111/Q111^* striatal cell lines ^27^ presenting endogenous levels of normal or mutant huntingtin (with 7 and 111 glutamines, respectively) were used. Cells were maintained at 33°C and 5% CO_2_ in DMEM (Sigma-Aldrich, ref. D5671) supplemented with 10% FBS, 1 mM sodium pyruvate, 2 mM L-glutamine, 1% penicillin-streptomycin and 400 µg/mL Geneticin (G418 Sulfate) (Thermo Scientific, ref. 11811-023). Transformed Mouse Embryonic Fibroblast (MEFs) lines derived from YAC128 mice ^7^ and zQ175 knock-in mice were kindly provided by Dr. Gillian Bates (Huntington’s Disease Centre and UK Dementia Research Institute at UCL). YAC128 MEFs carry wild-type mouse (*Mm*) *Htt* mRNA (7 CAG) and a full-length human (*Hs*) *HTT* transgene modified in exon 1 to have a 125 glutamine repeat expansion (composed primarily of CAG codons but also containing 9 interspersed CAA codons) ^7,28^. Transformed zQ175 MEFs were established as previously described ^7^. They carry an exon 1 from human *HTT* with highly expanded CAG repeat (∼190 CAG repeats) ^29^. MEFs were maintained in DMEM supplemented with 10% fetal bovine serum (FBS), 1 mM sodium pyruvate, 2 mM L-glutamine, and 1% penicillin-streptomycin in a humidified incubator at 37°C with 5 % CO. HD cells were seeded in 6- or 12-well plates (for qPCR experiments), in 24-well plates (for immunocytochemestry) at a density of 1.5 x 10^6^ cells/cm^2^. Pharmacological inhibition of METTL3 was performed with STM2457 ^30^ at 10 µM and 20 µM during 48h.

### Plasmid design

Site-specific manipulation of m6A levels at *Htt* intron1 mRNA was achieved with the programmable RNA editing system dCas13b-ALKBH5. RNA editor construct was designed as previously described with some modifications. Briefly, RNA-targeting catalytically inactive Type VI-B Cas13 enzyme from *Prevotella sp P5-125* (dPspCas13b) ^31^ was fused to the m6A-demethylase ALKBH5 at the C-terminus of dCas13b with a six amino-acid (GSGGGG) linker. Plasmid design was performed as described elsewhere with some modifications ^32^. The spacer (SP) sequences bound to the guide RNA (gRNA) were designed based on the intron 1 sequence of *Htt*, upstream of the first cryptic poly(A) site at 680 bp. Further evaluation of the SP sequences was performed using the OligoAnalyzer Tool (IDT) and Nucleotide BLAST (NCBI) to avoid mismatch at off-target locations. Sequences of the SP were as follows: 5’-TAG TTA AAC CAG GTT TTA AGC ATA GCC AGA −3’ (SP-gRNA1); 5’-ACT CCA GTG CCT TCG CCG TTC CCA GTT TGC −3’ (SP-gRNA 2) and 5’-AGC CTT GTT GGG GCC TGT CCT GAA TTC GAT-3’ (SP-gRNA 3). The resulting fusion protein with its corresponding SP-gRNA was cloned into the PX458 vector by GenScript. HA tag was included for detection of the fusion protein. Blasticidin S deaminase gene was also inserted in the construct to allow for the generation of stable cell lines. To control for the effects of transfection itself and for steric hinderance, two control plasmids were used: catalytically dead ALKBH5 (H204A) ^33^ fused to dCas13b containing the same spacer sequence and the dCas13b plasmid without the demethylase (GeneScript).

### Cell transfections

Plasmid transfection was performed using the Lipofectamine 3000 reagent (Invitrogen, ref. L3000-008) following manufacturer’s instructions. For six-well assays, cells were transfected with 2.5 µg/well of the corresponding plasmid (dCas13b-ALKBH5, dCas13b-ALKBH5 H204A and dCas13b-control). Forty-eight hours after transfection, cells were treated with the appropriate concentration of selection antibiotic (6 µg/mL Blasticidin for ST*Hdh^Q7/Q7^* cells, and 4 µg/mL for ST*Hdh^+/Q111^* cells) and medium with Blasticidin was changed every 2-3 days. Polyclonal cells that had integrated the plasmid of interest were expanded and seeded for Western Blot analysis and RNA extraction.

Locked Nucleic Acid-Antisense oligonucleotides (LNA-ASOs) were transfected with Lipofectamine 3000 at the dosages indicated in the figure legends. The LNA-ASO complementary to the CAG repeat (LNA-CTG) consisted of a 20-nt length oligonucleotide, CTGCTGCTGCTGCTGCTGCT, with LNA located every third T and a phosphorothioate-modified backbone. LNA-CTG and the control scrambled LNA-modified sequence (LNA-SCB) 5′-GTGTAACACGTCTATACGCCCA-3′ were obtained from Qiagen as previously described in Rue *et al* ^34^

### RNA isolation

RNA from the corresponding biological samples was extracted using the RNeasy Lipid Tissue Mini Kit (Qiagen, ref. 74804), following the instructions of the manufacturer. Briefly, the frozen tissue was placed in QIAzol (Qiagen) and homogenized using a 25G syringe. In the case of cell cultures, growth medium was discarded, cells were washed once with PBS and then collected from the plates by directly adding QIAzol reagent and homogenizing with a scraper. The purified RNA was eluted in nuclease-free H_2_O and the quantity and quality was measured using the Nanodrop 1000 spectrophotometer (Thermo Fisher Scientific). Total RNA was subjected to DNase treatment (Sigma-Aldrich, ref. AMPD1) according to manufacturer’s instructions. Samples were stored at −80°C until use.

### MeRIP-qPCR

Relative quantification of m6A levels of the genes of interest was performed through m6A-RNA immunoprecipitation (MeRIP) -qPCR as described elsewhere ^35^. Briefly, 3-4.5 µg total non-fragmented RNA was tumbled with anti-m6A antibody (Synaptic Systems, ref. 202 111) conjugated to Protein A Dynabeads (Invitrogen, ref. 10002D) in IP buffer (150 mM NaCl, 10 mM Tris-HCl, pH=7.5, 0.1% IGEPAL CA-630 in nuclease-free H_2_O) at 4°C. The immunoprecipitated RNA was subjected to two rounds of competitive elution with an m6A containing buffer (45 µL of 5X IP buffer, 75 µL of 20 mM m^6^A (Sigma-Aldrich, ref. M2780), 7 µL of RNase inhibitor and 98 µL nuclease-free water) and the eluted RNA was then concentrated using the RNeasy MinElute Cleanup kit (Qiagen, ref. 74204). The immunoprecipitated RNA was then reverse transcribed and used for qPCR. The fold enrichment was determined by calculating the Ct values of the MeRIP sample relative to the input sample.

### MazF-qPCR

For validation and stoichiometric quantification of m6A sites, we followed the protocol published by Garcia-Campos *et al* ^36^, with minor modifications. Total RNA was heat denatured and digested with 10-20 U of MazF enzyme (TakaRa, ref. 2415A) for 15 min at 37°C. RNA was then subjected to a cleanup protocol using the RNeasy MinElute Cleanup kit (Qiagen, ref. 74204), followed by RNA elution in water. Primer-probe sets for qPCR analysis were designed flanking a potential m6A motif that contains the “ACA” sequence (test motif) and a control primer-probe set (no ACA control) with no “ACA” site was designed in a nearby region of the motif of interest. Methylation levels were calculated based on the Ct values obtained from MazF-digested and non-digested samples for the test motif and the no ACA control (Methylation ratio = ((MazF digested test motif/ MazF digested no ACA motif)/(non-digested test motif/non-digested no ACA motif))).

### cDNA synthesis and Real-time quantitative PCR (qPCR) assays

45-500 ng of total, MeRIP or MazF-digested RNA were reverse transcribed using the High Capacity cDNA Reverse Transcription Kit (Applied Biosystems, ref. 4368814) according to manufacturer’s instructions. PrimeTime qPCR Assays were purchased from Integrated DNA Technologies (IDT) to measure genes of interest and housekeeping genes (Supplementary Tables 3 and 4). The qPCR reaction was performed on 96-well plates in a final volume of 12 μL using the Premix Ex Taq Probe based qPCR assay (Takara Biotechnology, ref. RR390A). All reactions were run in duplicate on a StepOnePlus Real-Time PCR System (Applied Biosystems) set to the following cycling program: 1 cycle 95°C for 30 sec; 40 cycles 95°C for 5 sec, 60°C for 20 sec. Relative enrichment was calculated using the ΔΔCt method, with actin-ß (mouse) expression serving as housekeeping genes.

To evaluate the performance of reverse transcription in the presence of bound LNA-ASOs, we performed PCR reactions using Hotstaq Plus DNA Polymerase (Qiagen ref:203603). PCR amplification was performed using exon 1 sequence-specific primers (Supplementary Table 5). PCR products were loaded and run on a 2 % agarose gel.

### mRNA stability

ST*Hdh^Q111/Q111^* cells were stably transfected with a dCas13b with gRNA2 without ALKBH5, dCas13b-A5 combined with NT-gRNA (control) and dCas13b-A5 combined with gRNA 2 constructs. After 24h cells were treated with actinomycin D (Act-D, Catalog #A9415, Sigma, USA) at 10 μg/ml for 2, 4, 6 and 8 h. Cells were collected with RNA isolated for real-time PCR.

### 3’RACE

Amplification of poly(A) mRNA was performed using the 3’RACE System for Rapid Amplification of cDNA Ends (Invitrogen, ref. 18373-019) according to the manufacturer’s instructions. First strand cDNA synthesis of mouse striatal MeRIP samples (IP and input control) was performed following the instructions for transcripts with high GC content. 100ng of the RNA of interest were reverse transcribed using the provided adapter primer (AP), followed by RNase H digestion to get rid of the template. The resulting cDNA was then amplified with a gene specific primer (GSP) for *Htt1α* generated by the first cryptic poly(A) site (GSP-pA1) or the second cryptic poly(A) site (GSP-pA2). All PCRs were performed using the Platinum II Hot-Start PCR Master Mix (Invitrogen, ref. 14000-013), containing 8 µL of the cDNA template, 0.8 µL of 10 µM primers, 8 µL of the Platinum GC Enhancer and 20 µL of the master mix. The thermocycler was set to the following cycling program:1 cycle 94°C for 3 min; 35 cycles 94°C for 15 sec, 60°C for 15 sec, 68°C for 1 min 30 sec; followed by cooling to 4°C. Sequences of the primers used for 3’RACE are provided in Supplementary Table 6. Samples were run on a 2% agarose gel, bands of interest were extracted with the GeneJET Gel extraction Kit (Thermo Scientific, ref. K0691), and DNA was reamplified using the Platinum II Hot-Start PCR Master Mix and the corresponding GSP primers to ensure an optimal amplification product for SANGER sequencing. The amplification product was subjected to a PCR cleanup protocol using the ExoSAP-IT Express PCR Product Cleanup kit (Applied Biosystems, ref. 75001), followed by quantification on a Nanodrop 1000 spectrophotometer (Thermo Fisher Scientific). SANGER sequencing was performed on a ABI3730XL DNA Analyzer (Genomics Core Facility, Universitat Pompeu Fabra).

### Immunocytochemistry

CRISPR-dCas13b stably transfected ST*Hdh^Q111/Q111^* immortalized striatal cells were grown on coverslips in 24-well plates for 48h and fixed for 10 min. at room temperature, permeabilized for 10 min with 0.5% saponin in PBS and blocked with 15% horse serum in PBS. The cells were incubated with anti-phospho-Histone H2AX (Ser139) (1:1000; Merck) or anti-ALKBH5 (1:2000; Sigma Aldrich) and secondary antibody (Cy3, 1:500; Jackson ImmunoResearch). Single images were acquired digitally using Leica Confocal SP5 with a ×40 oil-immersion objective. Percentage of nuclei with ɣ-H2AX foci and the average of ɣ-H2AX foci per cell were analyzed using the cell image analysis software CellProfiler. At least 20 images for each condition in 3 independent experiments were analyzed.

### Statistical analysis

Raw data were processed using Excel Microsoft Office and for further analysis transferred to Graphpad Prism version 8.0.2. Results are expressed as mean ± SEM. Normal distribution was assessed with the Saphiro-Wilk test. For statistical analysis, unpaired Student’s t-test (two-tailed) two or one-way ANOVA was performed, and the appropriate post-hoc tests were applied as indicated in the figure legends. A 95% confidence interval was used, considering differences statistically significant when p < 0.05.

## RESULTS

### m6A-methylation levels of *Htt1a* are significantly increased in the striatum of *Hdh^+/Q111^* mice from early disease stages

Our previous study revealed by methylated RNA immunoprecipitation sequencing (MeRIP-seq) analysis a significant enrichment of m6A in the 5’ proximal region (278 bp downstream of 5’ splice site in the mouse sequence) of *Htt* intron 1 in hippocampal samples from *Hdh^+/Q111^* mice ^22^. To extend our findings to the most affected brain region in HD, we performed a MeRIP followed by qPCR to analyze m6A methylation levels of *Htt* transcripts in the striatum of *Hdh^+/Q111^*mice at 2 and 8 months of age as well as in striatal ST*Hdh* cells. qPCR analysis with different primer-probe sets spanning *Htt* was performed (Fig. 1a and Supplementary Fig 1a). The employed assays included: probes before the first cryptic poly(A) site (I1-pA1) able to identify *Htt1a* transcripts terminated at both cryptic poly(A) sites; probes before the second cryptic poly(A) site (I1-pA2) that only detects those transcripts terminated at the second poly(A) site; probes at the 3’ end of intron 1 (I1-3’) to identify the incompletely spliced intron 1 sequences that have not terminated at cryptic poly(A) signals; probes spanning intron 3 to account for unspliced pre-mRNA; and finally probes spanning exons 36 and 37 to determine full length spliced *Htt* (FL-*Htt*). As has been described in other knock-in mouse models ^5,37^, qPCR analysis of input samples detected higher levels of intronic sequences generated at I1-pA1 and I1-pA2 in *Hdh^+/Q111^* mice when compared to WT mice at both ages (Fig. 1b,d) as well as in striatal ST*Hdh* cells (Supplementary Fig. 1a). The levels of intronic sequences I1-3′ were comparable to intron 3, therefore WT intronic sequences levels detected were indicative of pre-mRNA background or any level of incomplete splicing in WT mice might be below the levels of detection as previously described ^37^. The increase of intronic sequences I1-PA1 and I1-PA2 were accompanied by a significant decrease in FL-*Htt* compared to WT levels at 8 months (Fig. 1e). Similar reduction was observed in ST*Hdh^Q111/Q111^* cells (Supplementary Fig. 1b). Through qPCR analysis of the m6A-immunoprecipitated RNA we detected a significant enrichment of I1-pA1 transcripts in the striatum of 2-(Fig.1f) and 8-months-old *Hdh^+/Q111^* mice (Fig. 1g) with a significant increase of m6A levels in I1-pA2 detected only at 8 months. We also observed a significant enrichment of I1-pA1 and I1-pA2 transcripts in ST*Hdh^Q111/Q111^* cells (Supplementary Fig. 1c). The observed enrichment in *Hdh^+/Q111^*and ST*Hdh^Q111/Q111^* cells was specific to *Htt1*a since no significant differences were observed in FL-*Htt* (Fig 1f,g). These results suggest that transcripts produced by aberrant splicing are enriched in m6A.

**Figure 1.**
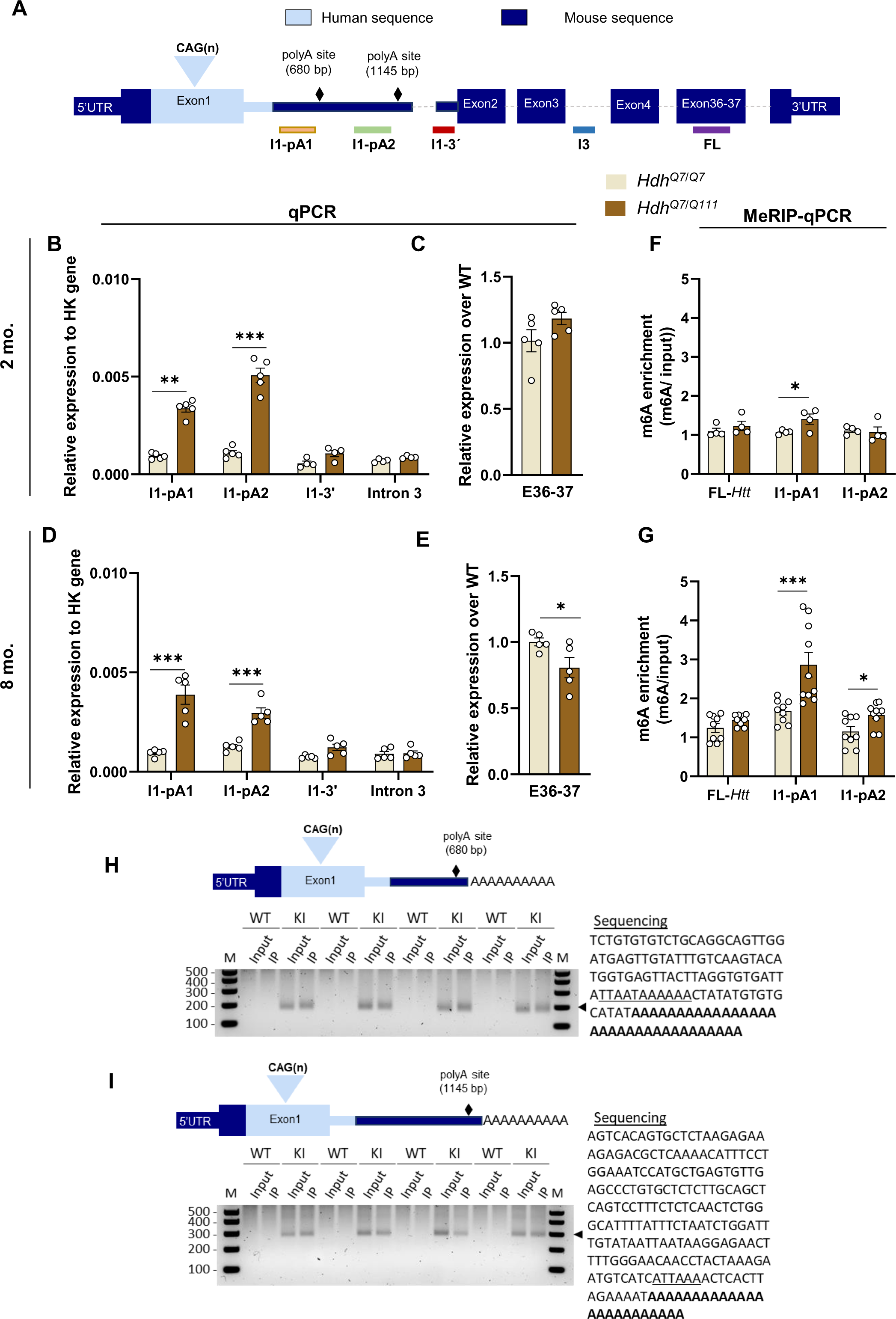
m6A methylation levels of *Htt1a* transcripts are increased in the *Hdh^+/Q111^* mouse model. **(A)** Schematic of the location of the primer-probe sets used for qPCR amplification of *Htt* transcripts. **(B**-**E)** qPCR analysis of *Htt* transcripts in the striatum of **(B,C)** 2 months (n=4-5 /genotype) and **(D,E)** 8 months old *Hdh*^+/Q111^ mice (n=5/genotype). Expression of intronic sequences is presented relative to housekeeping gene **(B,D)** and the relative levels of FL-*Htt* are shown relative to WT **(C,E). (F,G)** Analysis of m6A enrichment was measured by MeRIP-qPCR in the striatum of **(F)** 2 months, (n=4/ genotype) **(G)** 8 months old *Hdh*^+/Q111^ mice (n= 9-10/ genotype). Enrichment of m6A was measured by qPCR and normalized to input. **(H,I)** 3’RACE product in striatal samples generated from the cryptic poly(A) site at 680 bp **(H)** and 1145 bp **(I)** into intron 1 of *Htt*. (n=4 /genotype). M: DNA ladder. Right panel: SANGER sequencing of the generated product. The cryptic polyadenylation signal is underlined, and the poly(A) tail is shown in bold. Sequence was obtained from MeRIP and input samples (n=1 mouse for position 680 bp; n=2 mice for position 1145 bp). Data represent the mean ± SEM. Data were analyzed using Student’s two-tailed t-test. \**p* < 0.05 and \*\*\**p* < 0.001 compared with WT FL: full length; I: intron; pA: polyA site.

To confirm the m6A methylation in polyadenylated *Htt1a* transcripts generated by the cryptic poly(A) sites at 680 bp and 1,145 bp we performed 3’RACE of the input and the MeRIP RNA derived from striatal samples of 8 months old WT and *Hdh^+/Q111^* mice (Fig. 1h,i). Analysis of the input confirmed the presence of polyadenylated *Htt1a* generated by the two described cryptic poly(A) sites exclusively in *Hdh^+/Q111^* mice. Interestingly, when examining the MeRIP fraction, all *Hdh^+/Q111^* samples generated a 3’RACE product, indicating that polyadenylated mRNAs generated by aberrant splicing of *mHtt* were indeed m6A-modified.

### Aberrant m6A methylation at specific DRACH consensus motifs is conserved in human HD samples

Deposition of m6A mainly takes place at the DRACH consensus motif, where D=A, G or U; R= G or A; and H=A, C or U ^38–40^. Accurate identification of m6A sites in specific mRNAs is invaluable for better understanding their biological functions. Therefore, we further confirmed our results with an antibody-independent, single-base resolution method. We interrogated selected m6A sites in *Htt* intron 1 from different HD cell lines by MazF-qPCR, an approach that relies on the ability of the bacterial RNase MazF to cleave RNA at unmethylated sites occurring at ACA motifs, but not at the methylated counterparts m6A-CA ^41^. Potential m6A sites containing ACA motifs were chosen based on the location of the m6A peak previously detected as enriched in *Htt* RNA via MeRIP-seq (Supplementary Figure 2) ^22^. Those ACA sites were all located within the first 525 bp of *Htt* intron 1. We designed different sets of primers to amplify the regions containing those m6A consensus motifs, and we analyzed their methylation levels using MazF-qPCR analysis in three different HD cell lines (Fig 2a). We used the ST*Hdh^Q111/Q111^*cells which express the chimeric human/mouse (hm/ms) mutant *Htt* gene that contains 268 bp of human intron 1; embryonic mouse fibroblasts (MEFs) from zQ175 mice which express mutant *Htt* in which 84 bp of the 5′ end of mouse intron 1 have been replaced with 10 bp from the 5′ end of human intron 1 ^29^, and MEFs from YAC128 mice which express human mutant *HTT* ^7^. qPCR analysis in ST*Hdh^Q111/Q111^*cells revealed an increased methylation ratio in human GGACA and mouse AGACA compared with a position located further downstream in the mouse intron 1 in which significantly decreased levels of m6A levels were detected (GGACA ms) (Fig. 2b). On the other hand, we observed that in MEF zQ175 cells the AGACA mouse motif appears more methylated than the downstream GGACA mouse site (Fig. 2c). In YAC128 MEFs we also found significant differences between the different sites analyzed with the highest methylation ratio observed in the GGACA site (Fig. 2d) as we also observed in the ST*Hdh^Q111/Q111^* cells (Fig. 2b). These data suggest that the single peak of m6A in intron 1 reflects the overall levels of methylation of more than one m6A site. These data further suggest that methylation occurs at different rates in different co-existing m6A motifs favoring motifs closer to the 5’ of intron 1.

**Figure 2.**
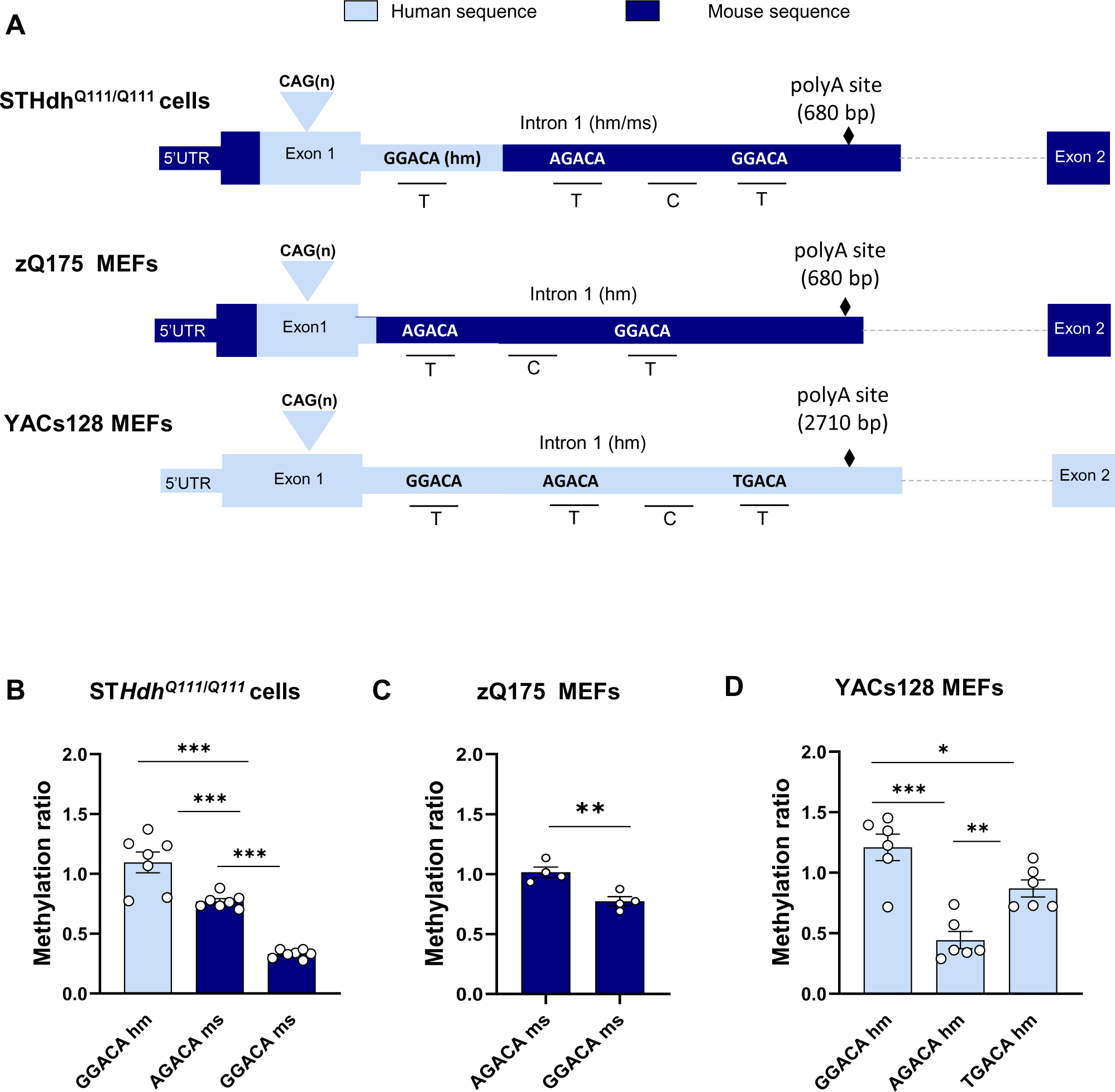
Interrogation of methylation levels at selected m6A DRACH motifs in the proximal region of m*Htt* intron 1. **(A)** Schematic of m*Htt* intron 1 m6A motifs analyzed by a qPCR-based assay coupled with MazF digestion in HD cell models, ST*Hdh^Q111/Q111^,* zQ175 MEFs and YAC128 MEFs cells **(B-D)** Methylation ratio obtained by MazF-qPCR analysis in **(B)** ST*Hdh^Q111/Q111^* cells (n=7 technical replicates/ motif), **(C)** zQ175 MEFs (n=4 technical replicates/motif) and **(D)** YAC128 MEFs (n=6 technical replicates/motif). The levels of a targeted amplicon (labeled ‘‘T’’) are measured against a control (labeled ‘‘C’’) amplicon in a MazF-digested sample and normalized against a nondigested sample. Data represent the mean ± SEM. Data were analyzed using One-way ANOVA with Tukey’s multiple comparisons test. \**p* < 0.05; \*\**p* < 0.01 and \*\*\**p* < 0.001. hm: human (light blue); ms: mouse (dark blue)

Next, we further evaluated the pathological relevance of m6A methylation in the 5’ region of m*Htt* intron 1 by analysis of the methylation ratio in HD *post-mortem* samples and HD skin-derived fibroblasts. We focused on the human GGACA site since was highly methylated in *Hdh^+/Q111^* and YAC128 MEFs cells expressing the human intronic sequence (Fig 3a). A gradual increase in m*HTT* intron 1 methylation was detected along progression of the neuropathology, showing significant differences in patients with Vonsattel grades 2-3 and 3 when compared to controls (Fig. 3b). Similarly, MazF-qPCR analysis in HD skin fibroblasts showed significant differences compared with controls (Fig. 3c) with increased methylation levels as disease progresses from pre-symptomatic to symptomatic stages. These data suggest that aberrant accumulation of m6A with disease progression could play a role in the worsening of HD symptomatology.

**Figure 3.**
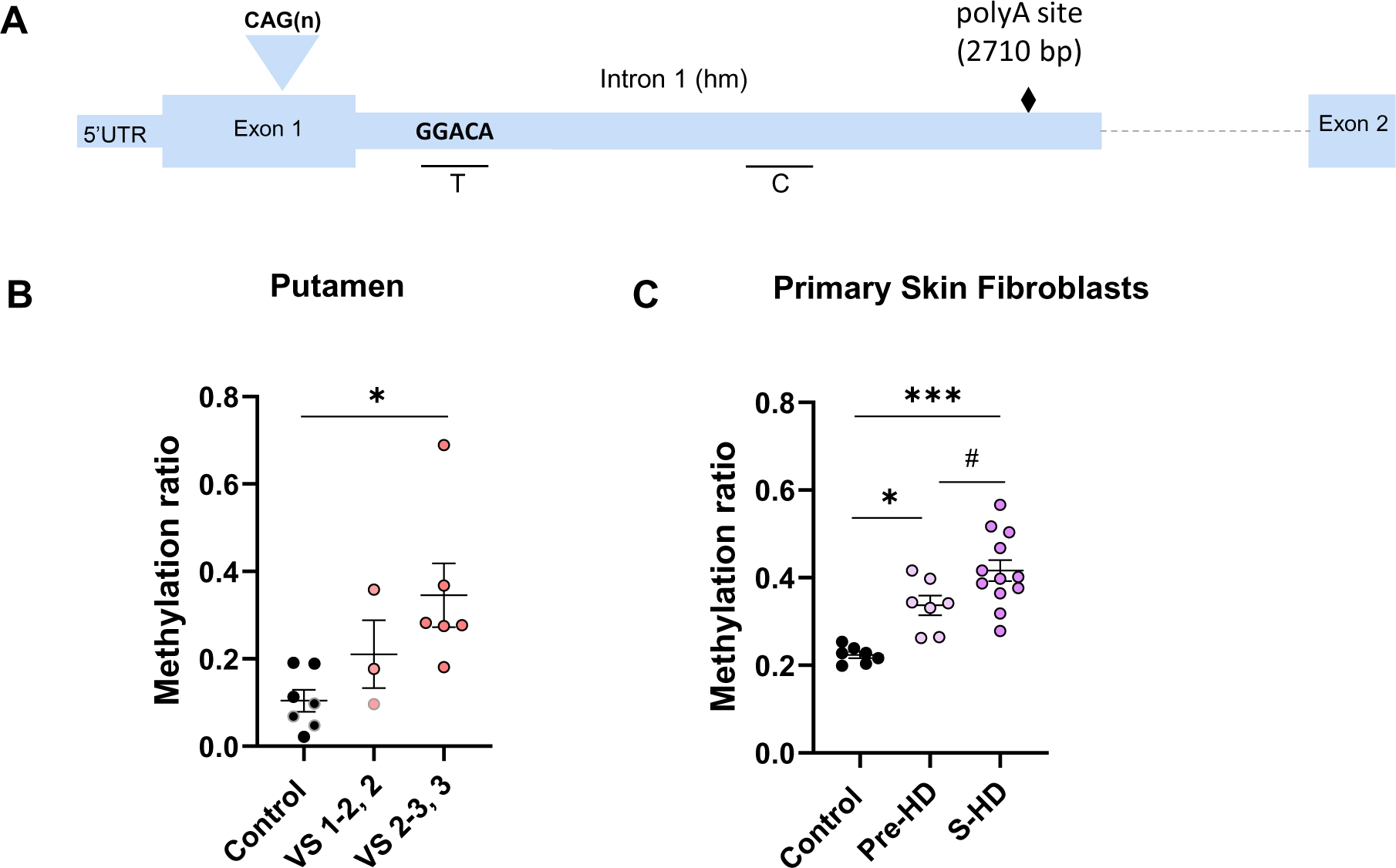
Increased methylation levels are detected at the m6A GGACA motif of *Htt* intron 1 in human samples. **(A)** Schematic of the DRACH motif GGACA hm in *HTT* intron 1 analyzed by a qPCR-based assay coupled with MazF digestion in human samples **(B-C)** Methylation ratio obtained by MazF-qPCR analysis in **(B)** human *post-mortem* samples of the putamen (n= 3-7 patients/group) and **(C)** in human skin fibroblasts (n=7-12 patients/group). VS: Vonsattel grade. Pre: presymptomatic; S: symptomatic. The levels of a targeted amplicon (labeled ‘‘T’’) are measured against a control (labeled ‘‘C’’) amplicon in a MazF-digested sample and normalized against a nondigested sample. Data represent the mean ± SEM. Data were analyzed using One-way ANOVA with Tukey’s multiple comparisons test. \**p* < 0.05; \*\*\**p* < 0.001 compared to control; #*p* < 0.05 compared to pre-HD. hm: human

### METTL3 modulates *Htt* intron1 methylation and reduces the expression levels of *Htt1α* transcripts

The METTL3/14 writer complex regulates the deposition of m6A in both intronic and exonic m6A regions ^42^ modulating several aspects of the mRNA lifecycle, including splicing ^23, 43^. To analyze the potential role of METTL3 in the methylation of *Htt* intron 1 and expression of the *Htt* transcripts we inhibited METTL3 with STM2457, a novel METTL3-specific inhibitor, that can bind to the S-adenosyl-L-methionine (SAM) binding site ^44^. qPCR and MazF-qPCR were performed in ST*Hdh* cells treated at 10 µM and 20 µM STM2547 for 48h. MazF-qPCR analysis of *Htt* intron 1 revealed a significant decrease of m6A levels in ST*Hdh^Q111/Q111^* cell in the GGACA site present in the human region of the chimeric m*Htt* intron 1 in response to the inhibition of METTL3 (Fig. 4a). When analyzing the levels of *Htt* FL and *Htt*1*a* intronic sequences before the first cryptic polyA site (I1-pA1) via qPCR, a significant reduction of only I1-pA1 could be detected in ST*Hdh^Q111/Q111^* cells at both concentrations of STM2457 when compared to the vehicle (Figure 4b), showing no changes in the relative levels of Intron 3 (Supplementary Fig. 3) or in the levels of FL-*Htt* and I1-pA1 sequences in ST*Hdh^Q7/Q7^* cells treated with STM2457 (Supplementary Fig.3). Similar results were observed when we analyzed methylation ratio and *HTT* transcripts in YAC128 MEFs. Inhibition of METTL3 decreased the ratio of methylation at the GGACA site without affecting the other motifs analyzed (Fig. 4c). Notably, this reduction of m6A levels was also accompanied by a significant decrease in *HTT1*a but not FL-*HTT* transcripts (Fig. 4d). We also evaluated the impact of METTL3 inhibition in zQ175 MEFs which do not contain the human intronic region with GGACA human motif. Consistent with our observations in ST*Hdh^Q111/Q111^* and YAC128 MEFs cells we did not find significant changes in the methylation levels when murine motifs (AGACA and GGACA) were analyzed (Fig. 4e). *Htt1a* levels were also downregulated by STM2457, albeit to comparatively lower levels than in ST*Hdh^Q111/Q111^* and YAC128 MEFs cells (Fig. 4f). These results ensured reliability of the human GGACA motif identified by the MazF-qPCR method and indicate that the methylation of this motif is dependent on METTL3 activity. Furthermore, a general reduction of m6A levels by METTL3 inhibition downregulates the expression of *Htt1a* suggesting a critical role of m6A modifications in *Htt / HTT* RNA metabolism.

**Figure 4.**
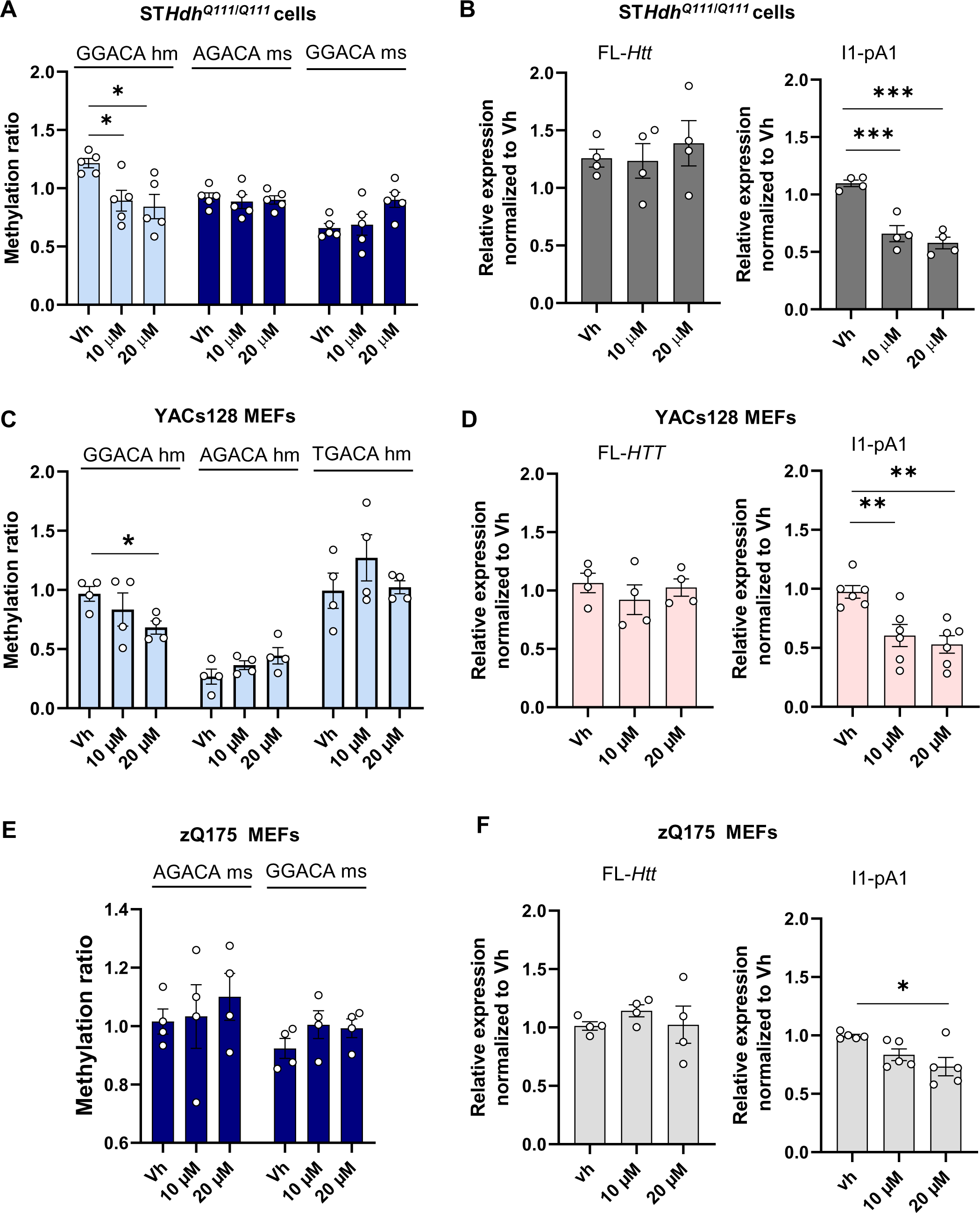
Pharmacological inhibition of METTL3 by STM2457 regulates the expression of *Htt1a* in HD *in vitro* models. **(A,C,E)** Methylation ratio obtained by MazF-qPCR analysis of *Htt* intron 1 in **(A)** ST*Hdh^Q111/Q111^* cells, **(C)** YAC128 and **(E)** zQ175 MEFs 7.1 treated with DMSO (Vh), STM2457 10 µM and STM2457 20 µM for 48h (n=8-9/ condition from 4 independent experiments). **(B,D,F)** qPCR analysis of *Htt* transcripts (FL-*Htt* and I1-pA1) in **(B)** ST*Hdh^Q111/Q111^* cells, **(D)** YAC128 and **(F)** MEFs 7.1 treated with DMSO (Vh), STM2457 10 µM and STM2457 20 µM for 48h. (n=5 independent experiments; 3 technical replicates/ experiment). Data represent the mean ± SEM. Data were analyzed using One-way ANOVA with Tukey’s multiple comparisons test. \**p* < 0.05; \*\**p* < 0.05; \*\*\**p* < 0.001 compared to vehicle (Vh).

### Demethylation of intron 1 using a CRISPR/dCas13b fused to Alkbh5 downregulates the expression of *Htt1a* transcripts

To elucidate causal relationships between the specific presence of m6A in intron 1 and the downregulation of *Htt1a* we conducted a targeted demethylation of *Htt* intron 1 in ST*Hdh^Q111/Q111^*cells using a CRISPR-Cas13-based approach. (Fig. 5a). We designed a plasmid construct that expresses the dCas13b-ALKBH5 fusion protein by linking the C-terminus of the inactive Cas13b (catalytically dead type VI-B Cas 13 enzyme named as dPspCas13b in Cox *et al.* ^31^) to the m6A demethylase ALKBH5 with a C-terminal HA tag. To target *Htt* intron 1, ST*Hdh* cells were stably transfected with three different constructs expressing the fusion protein and non-targeting gRNA (NTgRNA) or gRNAs designed to target three distinct positions, which locate inside intron 1 around the m6A sites previously described (gRNA1-2 and 3) (Supplementary Fig. 4). First, we analyzed the weight-shift of ALKBH5 indicating successful transfection of the cells (the fusion protein weighs around 150 kDa, which corresponds to the sum of dCas13 (124 kDa) and ALKBH5 (44 kDa) (Supplementary Fig. 5a). We then verified the effect of our RNA editing system by analyzing the methylation levels of m6A sites. MazF-qPCR showed that all three dCas13b-ALKBH5 systems expressing the different gRNAs significantly decreased the methylation ratio of the targeted GGACA human site compared with CTR (dCas13b-ALKBH5 NTgRNA) while no changes were observed in the other two motifs. The strongest and most significant demethylation was observed with gRNA2, which targets a 200 nt downstream region from the m6A GGACA site resulting in a 50 % demethylation (Fig. 5b). When analyzing the expression levels of the different *Htt* transcript isoforms, qPCR analysis showed that transfection with gRNA-2 and-3 significantly decreased the expression of transcripts generated by the first and second cryptic poly(A) sites while no differences were detected in FL-*Htt* levels (Fig. 5c). Next, we stably transfected cells with dCas13b-gRNA2 fused to H204A which is a catalytically inactive mutant of ALKBH5. Consistent with the previous results, stable transfection of ST*Hdh^Q111/Q111^* cells with the dCAs13b-ALKBH5 demethylase showed a significant reduction of about 20% of the *Htt1a* transcripts generated by the first and second cryptic poly(A) sites respectively. When cells were transfected with the catalytically inactive H204A enzyme, we found no difference, confirming that demethylation of *Htt* intron 1 RNA was achieved specifically by the RNA editing ALKBH5 constructs. In addition, consistent with previous observations, no significant changes were observed in FL-transcripts either when testing the probes against I1-3’ or the control I3 (Fig. 5f, Supplementary Fig. 5b). These results support a role of m6A in the expression of *Htt1a* transcripts.

**Figure 5.**
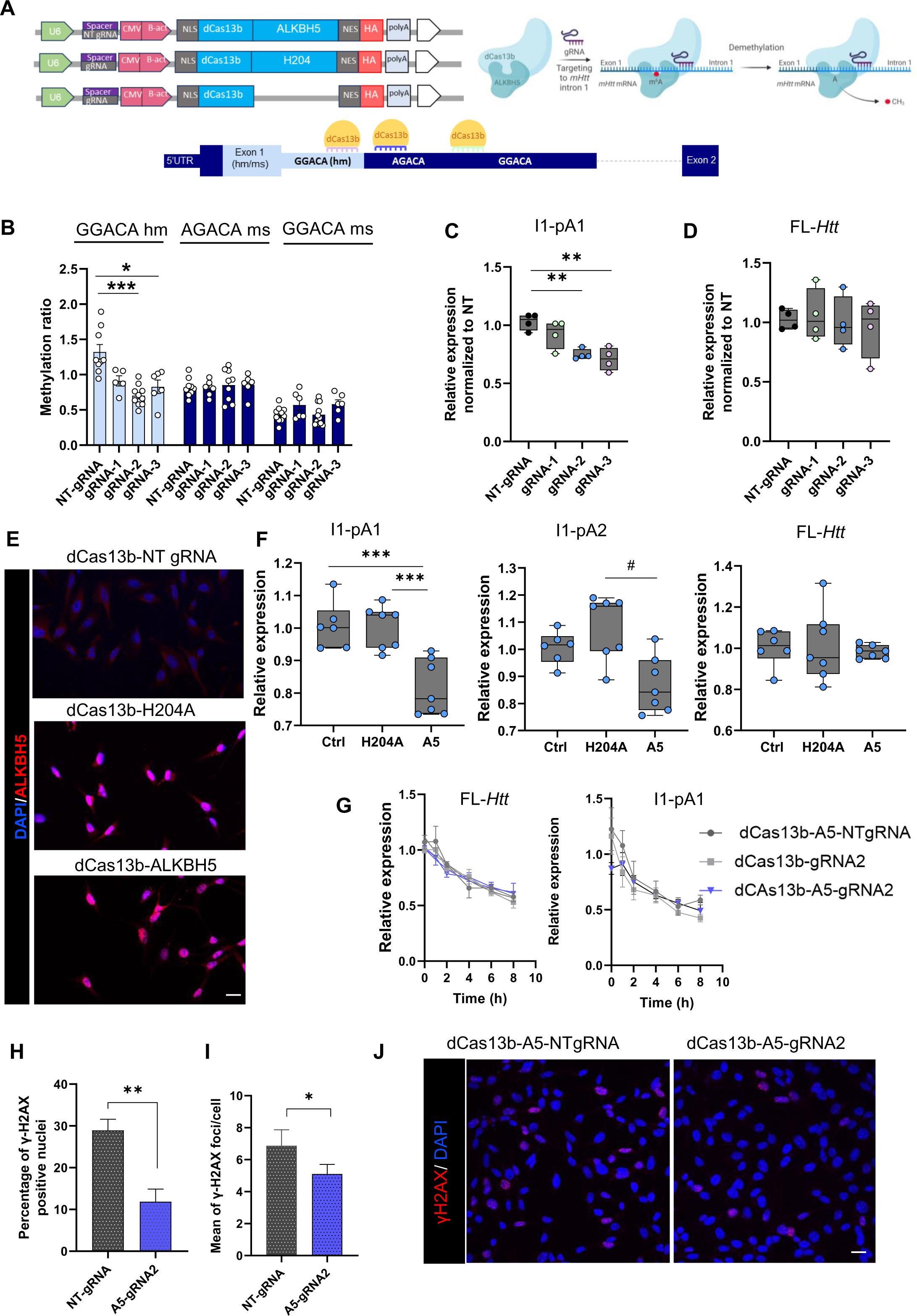
Target demethylation of *Htt* intron 1 regulates the expression of *Htt1a.* **(A)** Schematic representation of CRISPR dCas13b plasmid constructs and positions of m6A site within *Htt* intron 1 mRNAand regions targeted by three different gRNAs. **(B)** MazF-qPCR analysis of the DRACH motifs int *Htt* intron 1 (n=6-9 technical replicates from 3-4 independent experiments) and **(C-D)** qPCR analysis of expression levels of I1-pA1 **(C)** and FL-*Htt* transcripts **(D)** in ST*Hdh*^Q111/Q111^ cells transfected with dCas13b-ALKBH5 (dCas13b-A5) combined with NT-gRNA (control), gRNA 1, 2 and 3 (n=4 independent experiments; 3 technical replicates/ experiment). Data represent the mean ± SEM. Data were analyzed using One-way ANOVA with Tukey’s multiple comparisons test. \**p* < 0.05; \*\**p* < 0.05; \*\*\**p* < 0.001 compared to control (NT-gRNA). **(E)** Representative images from immunofluorescence staining of ALKBH5 in transfected ST*Hdh*^Q111/Q111^ cells with dCas13b (control, inactive H204A and ALKBH5). Nucleus is stained with DAPI (blue). **(F)** Expression levels of *Htt* transcripts (I1-pA1, I1-pA2, FL-*Htt*,) in transfected ST*Hdh*^Q111/Q111^ cells with dCas13b-NT gRNA, dCas13b-H204 gRNA2 and dCas13b-A5 gRNA2. (n=6-7 independent experiments) **(G)** RNA decay profile of *Htt* transcripts in transfected ST*Hdh*^Q111/Q111^ cells treated with Actinomycine-D (Act-D) for the indicated times (n=4 independent experiments; 2-3 technical replicates/ experiment) **(H-J)** Histograms showing the percentage of nuclei with ɣ-H2AX foci **(H)** and the average of ɣ-H2AX foci per cell **(I)** in transfected ST*Hdh*^Q111/Q111^ cells with dCAs13b-NT gRNA or dCAS13b-A5 gRNA2. Data were analyzed using Student’s t-test (10-15 images from 4 independent experiments). **(J)** Representative images of γ-H2AX foci (red) in ST*Hdh*^Q111/Q111^ cells. Nucleus is stained with DAPI (blue). Scale Bar, 20 µm.

To investigate whether dCas13b-ALKBH5-induced downregulation of *Htt1a* transcripts was caused by m6A-mediated mRNA decay, we performed RNA lifetime profiling by collecting and analyzing RNA of targeted demethylated and control samples obtained at different time points after transcription inhibition with actinomycin D (ActD) (Fig. 5g). RNA stability assay showed that targeted demethylation of intron 1 of m*Htt* with dCAS13b-ALKBH5 gRNA2 did not change the RNA half-life, displaying comparable RNA decay levels with control stable transfected cells (dCAs13b-ALKBH5-NTgRNA and dCAs13b-gRNA2) both when analyzing *Htt1a* and FL-*Htt*. These results indicate that demethylation in m*Htt* intron 1 does not influence the stability of *Htt1a* but rather affects other mechanisms such as the aberrant splicing thus regulating the production of *Htt1a* transcripts (Fig. 5g).

Given the evidence supporting persistent accumulation of DNA damage in neuronal DNA as one of the important contributing factors to early pathogenesis of HD ^45^ we further evaluated the potential impact of targeted *Htt1a* intron demethylation on basal DNA damage in the stably transfected cells. To this aim we analyzed the phosphorylation of histone H2AX in Serine 139 (gamma-H2AX), a marker of DNA double strand breaks, by immunocytochemistry. We detected a significant decrease in the percentage of nuclei with γH2AX foci and in the number of γH2AX foci per cell in cells stably transfected with dCAs13b-ALKBH5-gRNA2 compared with control transfected cells (Figure 5h-j). Together, our results show that m6A methylation in *Htt* intron1 is involved in the incomplete splicing that generates *Htt1a* and its regulation might influence DNA damage response and DNA repair mechanisms. It remains to clarify to what extent DNA damage is associated with altered levels of *HTT1a*.

### CAG repeat expansion regulate methylation in *Htt* intron 1 and affect the expression of *Htt1a*

Context -dependent features and RNA secondary structure play a key role in determining m6A deposition [44–46]. Therefore, we explored whether CAG-trinucleotide repeats which are known to form RNA stable hairpin structures with protein binding properties ^46–48^, could be contributing to m6A deposition in the proximal region of *Htt* intron 1. We used locked nucleic acid–modified antisense oligonucleotides complementary to the CAG repeat (LNA-CTG) that preferentially bind to mutant *Htt* to block CAG expansions ^34^. We transfected ST*Hdh^Q111/Q111^* cells with different concentrations of LNA-CTG or the analogous scrambled control LNA-ASO (LNA-SCB) and monitored LNA-CTG binding to the CAG repeat at exon 1 as well as *Htt* transcripts expression (FL-*Htt* and *Htt1a*). As previously demonstrated ^34^, the lack of *HTT* exon 1 (*HTT*-e1)1 mRNA amplification with primers spanning the CAG repeat supports LNA-CTG binding to the expanded transgene (Supplementary Fig. 6). PCR with primers amplifying Exon1 outside the CAG repeat (HTT-e1*) revealed no changes in HTT mRNA levels in accordance with published data ^34^. However, we detected by qPCR decreased levels of *Htt1a* with the highest concentration of LNA-CTG while no changes in FL-*Htt* were observed. (Fig. 6a). These data suggest that blocking expanded CAG repeats with LNA-CTG ASOs specifically affects the production of *Htt1a* favoring the concept that changes in expanded CAG structure and activity is crucial to produce *Htt1a* pathogenic species.

**Figure 6.**
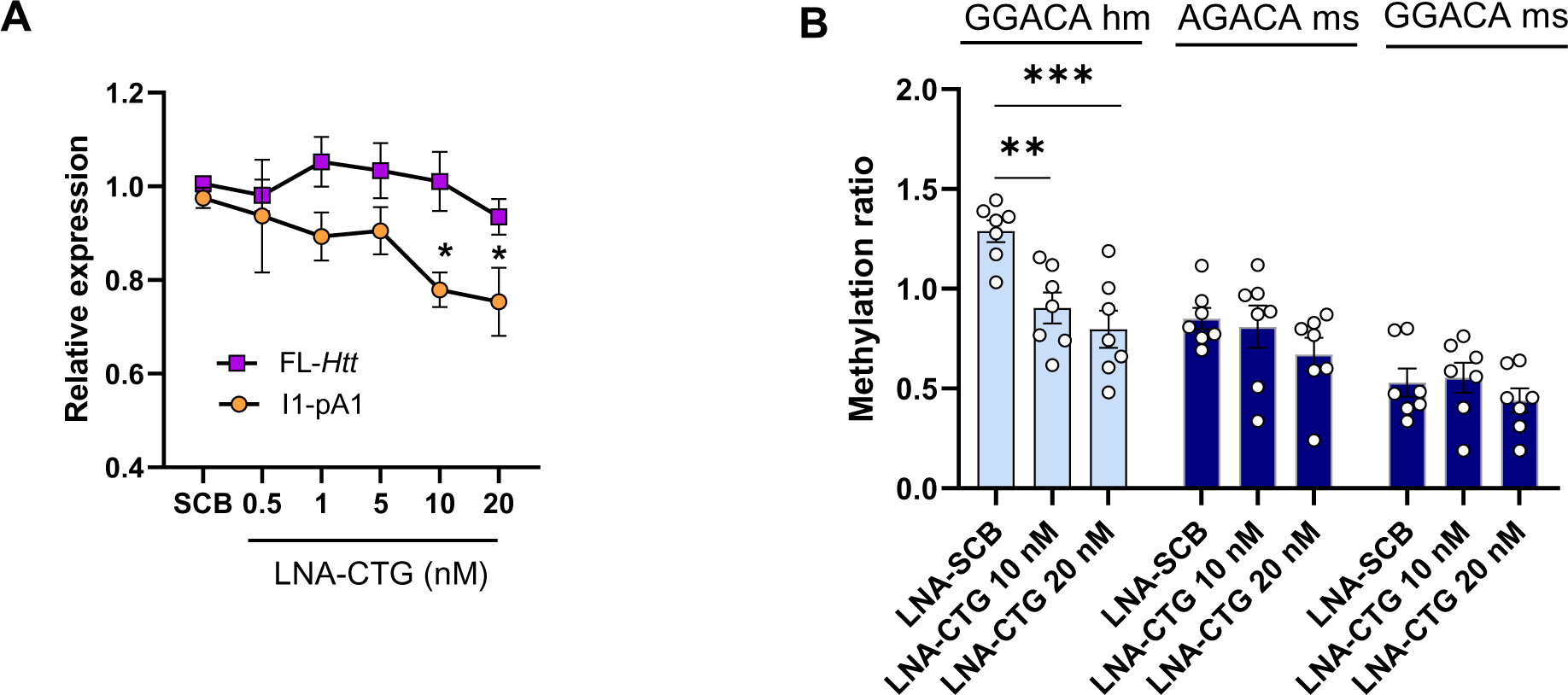
Blockage of expanded CAG repeats using LNA-CTG ASOs downregulates methylation in m*Htt* intron 1 RNA and decreases the levels of *Htt1a*. **(A)** qPCR analysis of expression levels of FL-*Htt* and I1-pA1 transcripts in ST*Hdh*^Q111/Q111^ cells transfected with LNA-SCB and LNA-CTG at 0.5, 1, 5, 10 and 20 nM (n= 3-8 independent experiments; 2 technical replicates/ experiment). Data represent the mean ± SEM. Data were analyzed using One-way ANOVA with Tukey’s multiple comparisons test. \**p* < 0.05 compared to LNA-SCB transfected cells. **(B)** MazF-qPCR analysis of methylation ratio of *Htt* intron 1 in ST*Hdh*^Q111/Q111^ cells transfected with 10 and 20 µM of LNA-CTG or LNA-SCB. Data represent the mean ± SEM. Data were analyzed using One-way ANOVA with Tukey’s multiple comparisons test. \*\**p* < 0.01, \*\*\**p* < 0.001 compared to LNA-SCB transfected cells.

Next, we determined the effect of blocking the activity of the CAG repeat expansion with the LNA-CTG on methylation levels of *Htt* intron 1 at the selected motifs here described. Methylation ratio obtained with MazF-qPCR analysis revealed a significant decrease in m6A abundance in the GGACA human motif of *Htt* intron 1 present in the ST*Hdh^Q111/Q111^* cells (Figure 6b). These results suggest that methylation of this specific site into intron 1 of m*Htt* RNA is directly affected by the CAG expansion, supporting the idea that expanded CAG plays a mechanistic role in modulating the axis *Htt* exon1-intron 1 methylation-*Htt1a* levels.

## DISCUSSION

Our study reveals that an aberrant m6A methylation in intron 1 of m*Htt* contributes to the generation of the mis-spliced mRNA variant, *Htt1a*. Building upon our previous evidence showing that *Htt* transcripts in the hippocampus of *Hdh^+/Q111^* mice are significantly methylated ^22^, our present results identify the m6A hypermethylation in *Htt* intron 1 in the striatum of *Hdh^+/Q111^*mice, as well as in HD cell lines and human samples. Inhibition of METTL3 and targeted demethylation of this intronic region regulate the expression of *Htt1a* suggesting that m6A contributes to the incomplete splicing of m*Htt*. This methylation is further influenced by CAG expansions. Collectively these data reveal a new CAG-dependent mechanism involved in the aberrant processing of m*Htt* relying on the m6A RNA modification.

The installation of m6A by the writer complex occurs co-transcriptionally, and its enrichment takes place in the last exon near the translation stop codon and 3′ untranslated region (3′ UTR) ^39, 40^. Recent studies have mapped m6A in the 5’UTR, exons and introns of nascent transcripts ^23, 42, 49, 50^, supporting earlier studies showing m6A on chromatin associated nascent pre-mRNA, including introns ^51, 52^. Here, we demonstrate that m6A is enriched in intronic sequences 5’ to the cryptic polyA sites expressed in *Htt1a* in the striatum of *Hdh^+/Q111^* mice and ST*Hdh^Q111/Q111^* cells. This finding is consistent with our MeRIP-seq data showing that *Htt* transcripts in the hippocampus of *Hdh^+/Q111^* mice presented a significant enrichment of m6A methylation in the 5’ proximal region of the intron. A similar enrichment has been reported toward the 5′-end of introns, particularly in regions involved in alternative splicing ^42, 50^. Therefore, our data suggest a potential functional role of m6A in alternative processing of *Htt* RNA.

Furthermore, we detected the m6A methylation in polyadenylated *Htt1a* mRNAs, indicating its persistence during RNA maturation and potential roles in *Htt1a* mRNA fate. For instance, it has been shown that *Htt1a* can be found in the nucleus in the form of RNA foci in YAC128 mice ^7^ and in HD *post-mortem* brains which are likely caused by somatic expansion of the CAG repeats ^8^. Although the direct function of m6A modifications in partitioning mRNAs into stress granules *in vivo* is unclear, it has been suggested that m6A, particularly in longer mRNAs containing multiple numbers of heavily modified m6A sites, can be involved in stress granule recruitment ^53^. It is intriguing to investigate whether m6A methylation, via interactions with scaffold reader proteins ^54–56^ contributes to *Htt1a* accumulation in nuclear clusters. This hypothesis is supported by a recent study showing that m6A is enriched in intronic polyadenylated (IPA) transcripts and that m6A regulates retention of mRNAs containing intact 5’splice site motifs in nuclear foci ^57^ m6A DRACH motifs are widespread throughout cellular transcriptomes, but only a small fraction was reported to be methylated *in vivo* ^39, 40^. Therefore, to cross-validate the known m6A sites within *Htt* intron 1, we studied the dynamic functions at a single-nucleotide resolution. We used an antibody-independent method to determine and quantify m6A methylation status present at specific sites in different HD cell lines. In our study we have identified the methylation of one motif at 147 nt from the exon1-intron1 splice junction in the human sequence of *STHdh^Q111/Q111^* and MEFs YAC128 cells which was consistently regulated by METTL3 inactivation or targeted demethylation. Of note, due to the limitations of the MazF-qPCR approach which can only quantify methylation levels at m6A motifs containing ACA sites [36,57], we cannot exclude methylation of other m6A sites which could not be assessed, particularly since the first intron of the huntingtin gene is highly enriched in DRACH consensus motifs[36,58]. Hence MEFs zQ175 cells which lack the human intronic region with the GGACA motif identified, could potentially have another m6A modified sites that were not analyzed in this work. Consistent with this, *Htt1a* levels decreased in zQ175 MEFs when METTL3 was inhibited. Further, our results in human samples demonstrate that m6A levels in the GGACA human motif increase with disease progression, suggesting that methylation at this site can be pathologically relevant.

The central question arising from the identification of m6A in this region of intron 1 is whether the modification is involved in the aberrant processing of m*HTT* RNA. Our results in HD cell lines show that inhibition of METTL3 activity resulted in a specific reduction of around 50% of *Htt1a* levels without changing expression levels of FL-*Htt*. Although this effect could be attributed to the observed STM2457-induced reduced levels of m6A in *Htt* intron 1, we cannot rule out the impact of reduced m6A modifications in other factors that could be critically involved in *Htt* RNA processing. For instance, on the one hand it has been shown that METTL3 regulates RNA splicing through m6A-mediated translational control of splicing factors ^58^. A broad distribution of m6A modifications across the CDS and 3’UTR has been demonstrated to regulate the expression of several targets of TDP43 ^59^, a nuclear RNA-binding protein integrally involved in RNA processing and previously associated with HD pathology ^60, 61^. Interestingly, a recent study proposes a co-regulatory role of TDP-43 with m6A modification in post-transcriptional RNA processing in HD (^62^ On the other hand, m6A can mediate mRNA degradation through the combined effects of the YTHDF readers on m6A target transcripts ^63^. Thus, we propose that METTL3 could influence *Htt1a* expression by regulating deposition of m6A in m*Htt* intron 1 as well as in other m6A-dependent transcripts with potential roles in splicing or stability. Moreover, we cannot exclude the interaction of m6A with other RNA bindind proteins such as TDP43 involved in alternative splicing.

To elucidate whether m6A deposition in *Htt* intron 1 es directly involved in *Htt1a* generation we interrogated the effect of m6A modifications using a CRISPR/dCas13b system fused to ALKBH5 to demethylate *Htt* intron 1 in a target-specific manner. Recent studies have reported the potential of CRISPR technology in the targeting of m6A modifications. A fusion protein linking inactive dCas13b to truncated METTL3 ^64^ or ALKBH5 ^65, 66^ allowed site-specific m6A incorporation or removal, respectively with low off-target effects. The N6-methyladenosine editing with CRISPR/dCas13 has already been successfully applied to several cancer models by targeting aberrant methylation of oncogenes ^32, 67^, hence constituting a promising approach to modulate m6A at specific transcripts. Notably, using this system in our immortalized mutant ST*Hdh^Q111/Q111^* cells, we detected a higher reduction in m6A methylation in comparison with the pharmacological inhibition of METTL3 showcasing the unique advantages of this CRISPR-based approach. In contrast with the rough inhibition of METTL3, the utility of a CRISPR/Cas system provides optimal targeting ability for the removal of m6A on specific sites avoiding global m6A regulation interference ^68^. This precision enhances the reliability of results when investigating the biological functions of m6A. Indeed, the precise regulation of m6A in *Htt* intron 1 was associated with a moderate but consistent reduction of ∼ 20% in *Htt1a* without affecting expression of FL-*Htt* indicating that intronic m6A modification is specifically involved in the regulation of *Htt1a* expression ^5^. The possibility that decreased levels of *Htt1a* observed by demethylation are a consequence of RNA degradation were ruled out by performing an RNA decay assay suggesting that m6A modifications may not affect the stability of *Htt1*a as previously described for other m6A-containing mRNAs ^69^ Thus, we conclude that our RNA editing system mediates efficient m6A demethylation in *Htt* intron 1 and allows to establish a causal relationship between m6A deposited in *Htt* RNA and *Htt1a* generation.

Several studies have documented splicing changes in the absence of an m6A writer ^70–72^, reader ^21, 73, 74^, or eraser ^75, 76^. Here we propose that one possible mechanism for m6A-dependent aberrant splicing regulation in m*Htt* is through pausing of Pol II ^21, 24^ and R-loop formation to facilitate transcription termination ^25^. Particularly, Pol II pausing, which can be regulated by m6A during transcription elongation, has been suggested to be modulated by RNA sequence and structure, chromatin state, and RNA-protein interactions ^77^. This is in line with evidence showing that several elements in intron 1 can reduce the elongation rate of Pol II, which could favor recognition of the cryptic poly(A) sites in intron 1 leading to generation of *HTT1a* ^4^. Therefore, our findings highlight intronic m6A modifications as a potential mechanism contributing to aberrant splicing of *mHtt*. Further investigations are required to elucidate whether m6A influences mis-splicing by directly controlling the pausing of RNA Pol II or promoting stalling of RNA Pol II by favoring R-loop formation.

To evaluate how m6A levels in *Htt* intron 1 impacts HD pathology we performed site-specific manipulation of m6A levels and DNA damage was evaluated in ST*Hdh^Q111/Q111^* cells which are known to show an augmentation of stress pathways, activated DNA damage response and apoptotic signals ^27,78^. Our results indicate that targeted demethylation of intron 1 reduces DNA damage as observed by fewer positive nuclei for γH2AX and formation of a smaller number of foci per nuclei. The N-terminus of HTT protein has been shown to disrupt DNA damage repair mechanism(s) leading to the excessive accumulation of DNA damage/strand breaks ^79^ and to localize in sites of DNA damage ^79^. Thus, the observed effect in our experiment could be driven by a reduction in the production of the toxic 90 aa N-terminal HTT-exon1 protein as a consequence of the downregulation of *Htt1a* transcripts. However, in line with the potential role of *Htt1*a in disease progression by formation of RNA foci at transcriptional sites ^8^, it is also possible that reduced methylation in *Htt1a* would reduce the interaction with other *Htt* transcripts in RNA clusters avoiding for instance, sequestering of RNA binding proteins involved in DNA damage ^80^.

Finally, we investigated whether expanded CAGs influence the deposition of m6A in this proximal site of intron 1. We used LNA-CTG ASOs that strongly bind to the CAGs RNAs, potentially disrupting their secondary structure and/or blocking their activity in exon 1 ^34^. Our data show that ASOs downregulated methylation levels in intron 1 and *Htt1a* levels suggesting that CAG expansions might be involved in this methylation. It has been demonstrated that m6A deposition can be determined by RNA secondary structure, by sequence motifs and by exon-intron architecture ^67, 81–83^. Indeed, it has been recently shown that the loss of endogenous RNA exon architecture and exon junction complexes (EJC) protection may result in m6A hypermethylation ^84^. In the context of *Htt* exon1, CAG-repeats form RNA stable hairpin structures ^47^ which aberrantly interact with several proteins, majority of them belonging to the spliceosome pathway ^11^. Thus, one possibility is that this secondary structure affects the dynamics of the Methyltransferase complex or that the recruitment of EJC proteins by the CAGs deprotect this proximal region resulting in higher deposition of m6A during pre-mRNA *Htt* transcription. This evidence points to existence of context-dependent pathological features in guiding m6A to *Htt* RNA.

Overall, our findings may lay the ground for the development of novel therapeutic strategies, since disease-modifying therapies targeting the pathological processing of *Htt* mRNA could be the most promising ^85^. While strategies involving the use of ASOs targeting HTT exon1-intron1 junction for lowering *Htt1a* or for the modulation of the aberrant splicing occurring in HD mutation carriers has been suggested ^86^, no evidences have been reported so far. When designing ASOs to target *Htt1a*, it might be relevant to avoid the m6A modified sites in *Htt* intron1 RNA since they could hinder ASO binding or lead to ASO destabilization due to improper base pairing, thereby reducing ASOs efficiency. Moreover, our evidence demonstrates that the CRISPR/dCas13b-ALKBH5 approach could allow a moderate reduction of the toxic mutant *HTT1a* fragment without affecting the WT allele. Intriguingly, a reduction by 43% of the *mHtt* mRNA has been reported to be enough in preventing further brain loss in symptomatic R6/2 mice and significantly increase lifespan using non-allele-specific ASOs, ^87^. In contrast, our strategy selectively targets the biogenesis of a highly pathogenic transcript derived from m*Htt*, obtaining a reduction of around 20% by merely modifying the methylation status of m*Htt* RNA. Given that slight reductions in *mHtt* mRNA levels are enough to ameliorate HD symptomatology in mouse models, future experiments validating the effects of *Htt1a* demethylation *in vivo* could provide encouraging results.

This work provides evidence of the role of dysregulation of m6A modifications as a potential pathogenic mechanism influencing m*HTT* RNA metabolism. Although additional work is needed to deepen our understanding of the function of m6A in m*HTT* intron 1, this study establishes the basis for new gene therapy strategies based on this RNA modification profile to target the mutant RNA allele or to modify the outcome of the splicing reaction that generates *HTT1A*.

## STATEMENTS & DECLARATIONS

### Author contributions

A.P carried out most experiments, set up the different methods used in this study, analyzed, interpreted results, contributed with manuscript preparation; I.R performed experiments, analyzed and interpreted results; K.S generated MEFs zQ175 cells; A.E performed experiments and analyzed results; A.S analyzed MeripSeq data; D.T supervised design of dCas13b-ALKBH5 constructs; S.G discussed experiments and data; G.B discussed experiments and data; U.V.O interpreted data and discussed experiments and data and E.M supervised ASOs experimental design, discussed experiments, analyzed, interpreted data and contributed with manuscript preparation; VB conceived, supervised the study, perform experiments, analyzed and interpreted results, and wrote the manuscript.

All authors read and approved the final manuscript.

### Fundings

This work was supported by Ministerio de Economía y Competitividad (PID2020-116474RB-100 to V.B. and RTI2018-094374-B100 to S.G.); Hereditary Disease Foundation grant to V.B.; Centro de Investigaciones Biomédicas en Red sobre Enfermedades Neurodegenerativas (CIBERNED) to S.G.; PhD grant program by La Generalitat de Catalunya (2018FI_B_00487) to A.P; PhD grant from PID2020-116474RB-100 (RE2021-097199) to I.R

### Competing interests

The authors have no relevant financial or non-financial interests to disclose.

## Acknowledgements

We are very grateful to Ana Lopez and Maria Teresa Muñoz for technical assistance, Dr Teresa Rodrigo and the staff of the animal care facility (Facultat de Psicologia Universitat de Barcelona). We thank members of our laboratory for helpful discussion.

## Data availability

Supplementary tables and information will be provided under reasonable request.

## Study approval

All procedures were performed in compliance with the National Institutes of Health Guide for the Care and Use of Laboratory Animals and approved by the local animal care committee of the Universitat de Barcelona (448/17) and the Generalitat de Catalunya (9878 P2), in accordance with the European (2010/63/EU) and Spanish (RD53/2013) guidelines for the care and use of laboratory animals.

## Notes

### Competing Interest Statement

The authors have declared no competing interest.

## REFERENCES

1. McColgan, P. & Tabrizi, S. J. Huntington’s disease: a clinical review. Eur J Neurol 25, 24– 34 (2018).

2. Walker, F. O. Huntington’s disease. Lancet 369, 218–228 (2007).

3. Ross, C. A. & Tabrizi, S. J. Huntington’s disease: from molecular pathogenesis to clinical treatment. Lancet Neurol 10, 83–98 (2011).

4. Neueder, A., Dumas, A. A., Benjamin, A. C. & Bates, G. P. Regulatory mechanisms of incomplete huntingtin mRNA splicing. Nat Commun 9, 3955 (2018).

5. Sathasivam, K. et al. Aberrant splicing of *HTT* generates the pathogenic exon 1 protein in Huntington disease. Proceedings of the National Academy of Sciences 110, 2366– 2370 (2013).

6. Mangiarini, L. et al. Exon 1 of the HD Gene with an Expanded CAG Repeat Is Sufficient to Cause a Progressive Neurological Phenotype in Transgenic Mice. Cell 87, 493–506 (1996).

7. Fienko, S. et al. Alternative processing of human *HTT* mRNA with implications for Huntington’s disease therapeutics. Brain 145, 4409–4424 (2022).

8. Ly, S. et al. Mutant huntingtin messenger RNA forms neuronal nuclear clusters in rodent and human brains. Brain Commun (2022) doi:10.1093/braincomms/fcac248.

9. Neueder, A. et al. The pathogenic exon 1 HTT protein is produced by incomplete splicing in Huntington’s disease patients. Sci Rep 7, 1307 (2017).

10. Smith, E. J. et al. Early detection of exon 1 huntingtin aggregation in zQ175 brains by molecular and histological approaches. Brain Commun 5, (2022).

11. Schilling, J. et al. Deregulated Splicing Is a Major Mechanism of RNA-Induced Toxicity in Huntington’s Disease. J Mol Biol 431, 1869–1877 (2019).

12. Gipson, T. A., Neueder, A., Wexler, N. S., Bates, G. P. & Housman, D. Aberrantly spliced *HTT*, a new player in Huntington’s disease pathogenesis. RNA Biol 10, 1647–1652 (2013).

13. Satterlee, J. S. et al. Novel RNA Modifications in the Nervous System: Form and Function. The Journal of Neuroscience 34, 15170–15177 (2014).

14. Cantara, W. A. et al. The RNA modification database, RNAMDB: 2011 update. Nucleic Acids Res 39, D195–D201 (2011).

15. Roundtree, I. A., Evans, M. E., Pan, T. & He, C. Dynamic RNA Modifications in Gene Expression Regulation. Cell 169, 1187–1200 (2017).

16. Shafik, A. M. et al. N6-methyladenosine dynamics in neurodevelopment and aging, and its potential role in Alzheimer’s disease. Genome Biol 22, 17 (2021).

17. Zhao, F. et al. METTL3-dependent RNA m6A dysregulation contributes to neurodegeneration in Alzheimer’s disease through aberrant cell cycle events. Mol Neurodegener 16, 70 (2021).

18. Xiao, W. et al. Nuclear m 6 A Reader YTHDC1 Regulates mRNA Splicing. Mol Cell 61, 507–519 (2016).

19. Wang, S. et al. Dynamic regulation and functions of mRNA m6A modification. Cancer Cell Int 22, 48 (2022).

20. Roundtree, I. A. et al. YTHDC1 mediates nuclear export of N6-methyladenosine methylated mRNAs. Elife 6, (2017).

21. Zhou, K. I. et al. Regulation of Co-transcriptional Pre-mRNA Splicing by m6A through the Low-Complexity Protein hnRNPG. Mol Cell 76, 70–81.e9 (2019).

22. Pupak, A. et al. Altered m6A RNA methylation contributes to hippocampal memory deficits in Huntington’s disease mice. Cellular and Molecular Life Sciences 79, 416 (2022).

23. Louloupi, A., Ntini, E., Conrad, T. & Ørom, U. A. V. Transient N-6-Methyladenosine Transcriptome Sequencing Reveals a Regulatory Role of m6A in Splicing Efficiency. Cell Rep 23, 3429–3437 (2018).

24. Akhtar, J. et al. m6A RNA methylation regulates promoter-proximal pausing of RNA polymerase II. Mol Cell 81, 3356–3367.e6 (2021).

25. Yang, X. et al. m6A promotes R-loop formation to facilitate transcription termination. Cell Res 29, 1035–1038 (2019).

26. Wheeler, V. Length-dependent gametic CAG repeat instability in the Huntington’s disease knock-in mouse. Hum Mol Genet 8, 115–122 (1999).

27. Trettel, F. Dominant phenotypes produced by the HD mutation in STHdhQ111 striatal cells. Hum Mol Genet 9, 2799–2809 (2000).

28. Pouladi, M. A. et al. Marked differences in neurochemistry and aggregates despite similar behavioural and neuropathological features of Huntington disease in the full-length BACHD and YAC128 mice. Hum Mol Genet 21, 2219–2232 (2012).

29. Mason, M. A. et al. Silencing Srsf6 does not modulate incomplete splicing of the huntingtin gene in Huntington’s disease models. Sci Rep 10, 14057 (2020).

30. Yankova, E. et al. Small-molecule inhibition of METTL3 as a strategy against myeloid leukaemia. Nature 593, 597–601 (2021).

31. Cox, D. B. T. et al. RNA editing with CRISPR-Cas13. Science (1979) 358, 1019–1027 (2017).

32. Li, J. et al. Targeted mRNA demethylation using an engineered dCas13b-ALKBH5 fusion protein. Nucleic Acids Res 48, 5684–5694 (2020).

33. Feng, C. et al. Crystal Structures of the Human RNA Demethylase Alkbh5 Reveal Basis for Substrate Recognition. Journal of Biological Chemistry 289, 11571–11583 (2014).

34. Rué, L. et al. Targeting CAG repeat RNAs reduces Huntington’s disease phenotype independently of huntingtin levels. Journal of Clinical Investigation 126, 4319–4330 (2016).

35. Pupak, A. et al. Altered m6A RNA methylation contributes to hippocampal memory deficits in Huntington’s disease mice. Cellular and Molecular Life Sciences 79, 416 (2022).

36. Garcia-Campos, M. A. et al. Deciphering the “m6A Code” via Antibody-Independent Quantitative Profiling. Cell 178, 731–747.e16 (2019).

37. Papadopoulou, A. S. et al. Extensive Expression Analysis of Htt Transcripts in Brain Regions from the zQ175 HD Mouse Model Using a QuantiGene Multiplex Assay. Sci Rep 9, 16137 (2019).

38. Schwartz, S. et al. Perturbation of m6A Writers Reveals Two Distinct Classes of mRNA Methylation at Internal and 5′ Sites. Cell Rep 8, 284–296 (2014).

39. Meyer, K. D. et al. Comprehensive Analysis of mRNA Methylation Reveals Enrichment in 3′ UTRs and near Stop Codons. Cell 149, 1635–1646 (2012).

40. Dominissini, D. et al. Topology of the human and mouse m6A RNA methylomes revealed by m6A-seq. Nature 485, 201–206 (2012).

41. Garcia-Campos, M. A. et al. Deciphering the ‘m6A Code’ via Antibody-Independent Quantitative Profiling. Cell 178, 731–747.e16 (2019).

42. Wei, G. et al. Acute depletion of METTL3 implicates *N* ^6^ -methyladenosine in alternative intron/exon inclusion in the nascent transcriptome. Genome Res 31, 1395–1408 (2021).

43. Adhikari, S., Xiao, W., Zhao, Y.-L. & Yang, Y.-G. m ^6^ A: Signaling for mRNA splicing. RNA Biol 13, 756–759 (2016).

44. Yankova, E. et al. Small-molecule inhibition of METTL3 as a strategy against myeloid leukaemia. Nature 593, 597–601 (2021).

45. Pradhan, S. et al. Polyglutamine Expansion in Huntingtin and Mechanism of DNA Damage Repair Defects in Huntington’s Disease. Front Cell Neurosci 16, (2022).

46. Krzyzosiak, W. J. et al. Triplet repeat RNA structure and its role as pathogenic agent and therapeutic target. Nucleic Acids Res 40, 11–26 (2012).

47. Jasinska, A. Structures of trinucleotide repeats in human transcripts and their functional implications. Nucleic Acids Res 31, 5463–5468 (2003).

48. Galka-Marciniak, P., Urbanek, M. O. & Krzyzosiak, W. J. Triplet repeats in transcripts: structural insights into RNA toxicity. bchm 393, 1299–1315 (2012).

49. Ke, S. et al. m ^6^ A mRNA modifications are deposited in nascent pre-mRNA and are not required for splicing but do specify cytoplasmic turnover. Genes Dev 31, 990–1006 (2017).

50. Hu, L. et al. m6A RNA modifications are measured at single-base resolution across the mammalian transcriptome. Nat Biotechnol 40, 1210–1219 (2022).

51. Carroll, S. M., Narayan, P. & Rottman, F. M. N ^6^ -Methyladenosine Residues in an Intron-Specific Region of Prolactin Pre-mRNA. Mol Cell Biol 10, 4456–4465 (1990).

52. Salditt-Georgieff, M. et al. Methyl labeling of HeLa cell hnRNA: a comparison with mRNA. Cell 7, 227–237 (1976).

53. Khong, A., Matheny, T., Huynh, T. N., Babl, V. & Parker, R. Limited effects of m6A modification on mRNA partitioning into stress granules. Nat Commun 13, 3735 (2022).

54. Ries, R. J. et al. m6A enhances the phase separation potential of mRNA. Nature 571, 424–428 (2019).

55. Gao, Y. et al. Multivalent m6A motifs promote phase separation of YTHDF proteins. Cell Res 29, 767–769 (2019).

56. Fu, Y. & Zhuang, X. m6A-binding YTHDF proteins promote stress granule formation. Nat Chem Biol 16, 955–963 (2020).

57. Eliza S. Lee et al. N-6-methyladenosine (m6A) Promotes the Nuclear Retention of mRNAs with Intact 5’ Splice Site Motifs. 10.1101/2023.06.20.545713.

58. Wu, Y. et al. METTL3-Mediated m6A Modification Controls Splicing Factor Abundance and Contributes to Aggressive CLL. Blood Cancer Discov 4, 228–245 (2023).

59. McMillan, M. et al. RNA methylation influences TDP43 binding and disease pathogenesis in models of amyotrophic lateral sclerosis and frontotemporal dementia. Mol Cell 83, 219–236.e7 (2023).

60. Sanchez, I. I. et al. Huntington’s disease mice and human brain tissue exhibit increased G3BP1 granules and TDP43 mislocalization. Journal of Clinical Investigation 131, (2021).

61. Tada, M. et al. Coexistence of Huntington’s disease and amyotrophic lateral sclerosis: a clinicopathologic study. Acta Neuropathol 124, 749–760 (2012).

62. Thai B. Nguyen et al. Aberrant splicing in Huntington’s disease via disrupted TDP-43 activity accompanied by altered m6A RNA modification. doi: 10.1101/2023.10.31.565004 (2023) doi:doi: 10.1101/2023.10.31.565004.

63. Zaccara, S. & Jaffrey, S. R. A Unified Model for the Function of YTHDF Proteins in Regulating m6A-Modified mRNA. Cell 181, 1582–1595.e18 (2020).

64. Wilson, C., Chen, P. J., Miao, Z. & Liu, D. R. Programmable m6A modification of cellular RNAs with a Cas13-directed methyltransferase. Nat Biotechnol 38, 1431–1440 (2020).

65. Li, J. et al. Targeted mRNA demethylation using an engineered dCas13b-ALKBH5 fusion protein. Nucleic Acids Res 48, 5684–5694 (2020).

66. Chen, X. et al. Targeted RNA *N* ^6^ -Methyladenosine Demethylation Controls Cell Fate Transition in Human Pluripotent Stem Cells. Advanced Science 8, (2021).

67. Gao, J., Luo, T., Lin, N., Zhang, S. & Wang, J. A New Tool for CRISPR-Cas13a-Based Cancer Gene Therapy. Mol Ther Oncolytics 19, 79–92 (2020).

68. Zhang, W., Qian, Y. & Jia, G. The detection and functions of RNA modification m6A based on m6A writers and erasers. Journal of Biological Chemistry 297, 100973 (2021).

69. Wang, X. et al. N6-methyladenosine-dependent regulation of messenger RNA stability. Nature 505, 117–120 (2014).

70. Haussmann, I. U. et al. m6A potentiates Sxl alternative pre-mRNA splicing for robust Drosophila sex determination. Nature 540, 301–304 (2016).

71. Lence, T. et al. m6A modulates neuronal functions and sex determination in Drosophila. Nature 540, 242–247 (2016).

72. Mendel, M. et al. Splice site m6A methylation prevents binding of U2AF35 to inhibit RNA splicing. Cell 184, 3125–3142.e25 (2021).

73. Kasowitz, S. D. et al. Nuclear m6A reader YTHDC1 regulates alternative polyadenylation and splicing during mouse oocyte development. PLoS Genet 14, e1007412 (2018).

74. Xiao, W. et al. Nuclear m 6 A Reader YTHDC1 Regulates mRNA Splicing. Mol Cell 61, 507–519 (2016).

75. Bartosovic, M. et al. N6-methyladenosine demethylase FTO targets pre-mRNAs and regulates alternative splicing and 3′-end processing. Nucleic Acids Res 45, 11356–11370 (2017).

76. Zhao, X. et al. FTO-dependent demethylation of N6-methyladenosine regulates mRNA splicing and is required for adipogenesis. Cell Res 24, 1403–1419 (2014).

77. Mayer, A., Landry, H. M. & Churchman, L. S. Pause & go: from the discovery of RNA polymerase pausing to its functional implications. Curr Opin Cell Biol 46, 72–80 (2017).

78. Cattaneo, M. et al. Longevity-Associated Variant of BPIFB4 Confers Neuroprotection in the STHdh Cell Model of Huntington Disease. Int J Mol Sci 23, 15313 (2022).

79. Gao, R. et al. Mutant huntingtin impairs PNKP and ATXN3, disrupting DNA repair and transcription. Elife 8, (2019).

80. Fijen, C. & Rothenberg, E. The evolving complexity of DNA damage foci: RNA, condensates and chromatin in DNA double-strand break repair. DNA Repair (Amst) 105, 103170 (2021).

81. Meiser, N., Mench, N. & Hengesbach, M. RNA secondary structure dependence in METTL3–METTL14 mRNA methylation is modulated by the N-terminal domain of METTL3. Biol Chem 402, 89–98 (2020).

82. Schwartz, S. et al. High-Resolution Mapping Reveals a Conserved, Widespread, Dynamic mRNA Methylation Program in Yeast Meiosis. Cell 155, 1409–1421 (2013).

83. Uzonyi, A. et al. Exclusion of m6A from splice-site proximal regions by the exon junction complex dictates m6A topologies and mRNA stability. Mol Cell 83, 237–251.e7 (2023).

84. He, P. C. et al. Exon architecture controls mRNA m ^6^ A suppression and gene expression. Science (1979) 379, 677–682 (2023).

85. Marxreiter, F., Stemick, J. & Kohl, Z. Huntingtin Lowering Strategies. Int J Mol Sci 21, 2146 (2020).

86. Tabrizi, S. J. et al. Targeting Huntingtin Expression in Patients with Huntington’s Disease. New England Journal of Medicine 380, 2307–2316 (2019).

87. Kordasiewicz, H. B. et al. Sustained Therapeutic Reversal of Huntington’s Disease by Transient Repression of Huntingtin Synthesis. Neuron 74, 1031–1044 (2012).

