## Supplemental Figures for "m6A RNA modification of m*Htt* intron 1 regulates the generation of *Htt1a* in Huntington’s Disease"

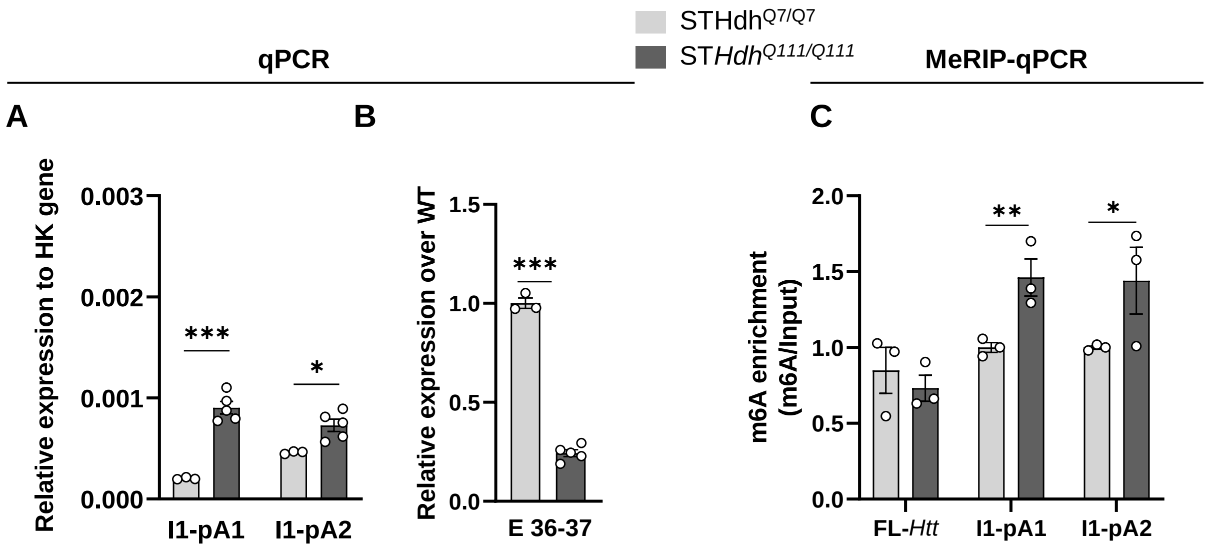


**Supplementary Figure 1. m6A methylation levels of *Htt1*a transcripts are increased in ST*Hdh*$^{Q111/Q111}$ cells.** qPCR analysis of **(A)** intronic sequences and **(B)** FL-*Htt* expression levels in the ST*Hdh* immortalized striatal cells (n=3-5/ genotype)**.** Expression of intronic sequences is presented relative to housekeeping gene and FL-*Htt* levels are shown relative to WT **(C)** MeRIP-qPCR analysis in the ST*Hdh* immortalized striatal cells (n= 3/4 genotype). Immunoprecipitated m6A transcripts are normalized to input and m6A enrichment is shown relative to WT. Data represent the mean ± SEM. Data were analyzed using Student´s two-tailed t-test. *$p < 0.05$, ***$p < 0.01$ and ***$p < 0.001$ compared with WT.

A

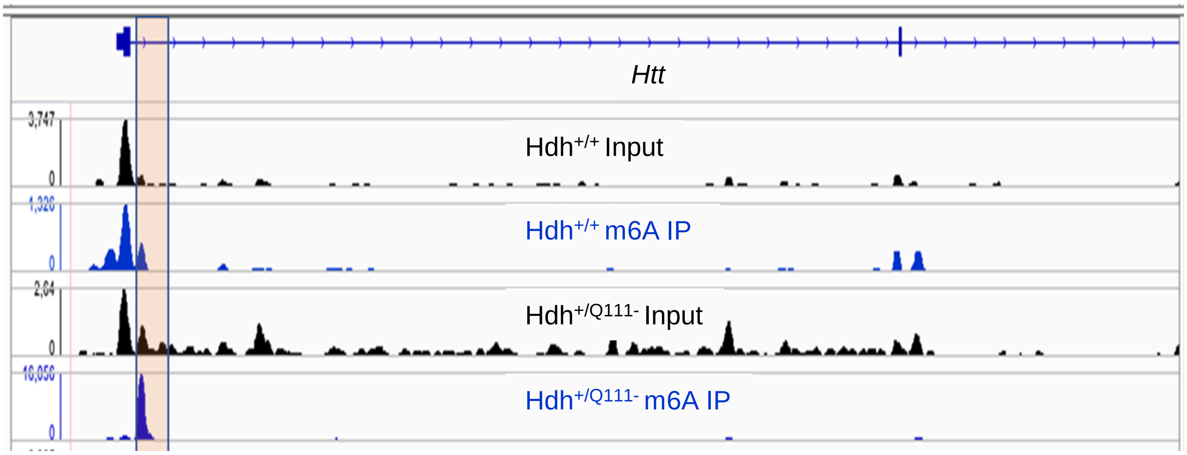


B

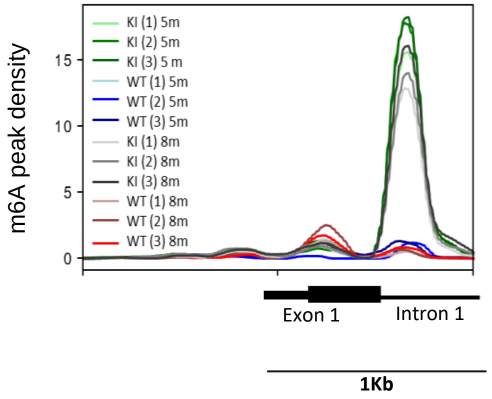


**Supplementary Figure 2: m6A enrichment in *Htt* intron 1 of *Hdh*$^{+/Q111}$ mice. (A)** Genome browser snapshots harboring m6A enrichment in the proximal region of *Htt* intron 1 to the 5' exon1-intron1 splice site. **(B)** Comparison of fold enrichment distribution of methylation sites in the *Htt* intron 1 between 8- and 5-month old WT and KI mice obtained by MeRIP-seq (n=3/ genotype/age, Pupak et al 2022)

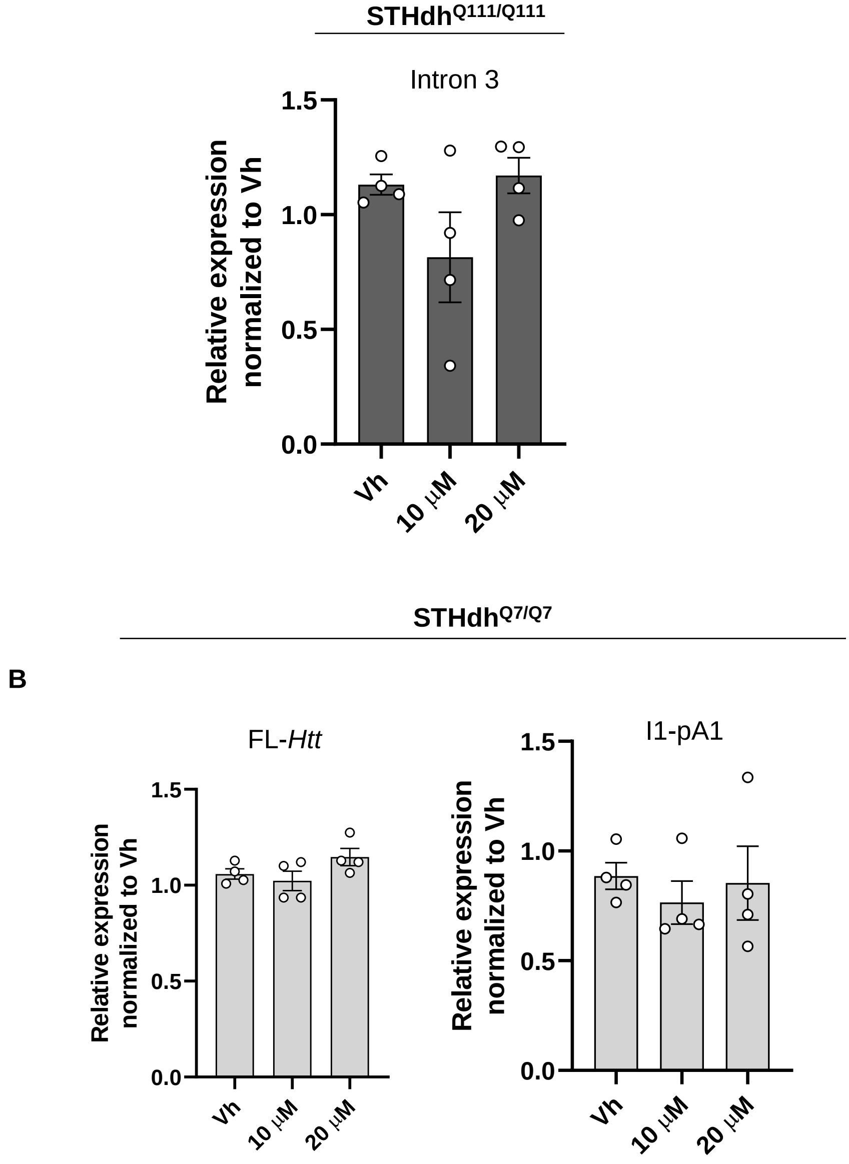


**Supplementary Figure 3. Pharmacological inhibition of METTL3 by STM2457 in ST*Hdh*$^{Q111/Q111}$ cells and ST*Hdh*$^{Q7/Q7}$ cells. (A)** qPCR analysis of the I3 *Htt* transcript in ST*Hdh*$^{Q111/Q111}$ cells treated with DMSO (Vh), STM2457 10 µM and STM2457 20 µM for 48h (n = 4 independent experiments). Data represent the mean ± SEM. Data were analyzed using One-way ANOVA with Tukey´s multiple comparisons test **(B)** qPCR analysis of FL-*Htt* and I1-pA1 transcripts in ST*Hdh*$^{Q7/Q7}$ cells treated with DMSO (Vh), STM2457 10 µM and STM2457 20 µM for 48h (n = 4 independent experiments). Data represent the mean ± SEM. Data were analyzed using One-way ANOVA with Tukey´s multiple comparisons test.

**A**

### Sequence hm/ms Exon1-Intron 1 of *Htt RNA in* ST*Hdh*$^{Q111/Q111}$ *cells and Hdh*$^{+/Q111}$ *mice*

GCACTCGCCGCGAGGGTTGCCGGGACGGGCCCAAGATGGCTGAGCGCCTTGGTTCCGCTTCTGCCTGCCGCGCAGAGCCCCAT
TCATTGCCTTGCTGCTAAGTGGCGCCGCGTAGTGCCAGTAGGCTCCAAGTCTTCAGGGTCTGTCCCATCGGGCAGGAAGCCGTC
ATGGCAACCCTGGAAAAGCTGATGAAGGCTTGCGGCGCCGCGAGTCGGCCCGAGGCCTCCGGGGACTGCCGTGCCGGGCGGG
AGACCGCCATGGCGACCCTGGAAAAGCTGATGAAGGCCTTCGAGTCCCTCAAGTCCTTCCAGCAGCAGCAGCAGCAGCAGCAG
CAGCAGCAGCAGCAGCAGCAGCAGCAGCAGCAGCAGCAGCAGCAGCAGCAGCAGCAGCAGCAGCAGCAGCAGCAGCAGCAG
CAGCAGCAGCAGCAGCAGCAGCAGCAGCAGCAGCAGCAGCAGCAGCAGCAGCAGCAGCAGCAGCAGCAGCAGCAGCAGCAG
CAGCAGCAGCAGCAGCAGCAGCAGCAGCAGCAGCAGCAGCAGCAACAGCCGCCACCGCCGCCGCCCCGCCGCCGCCTCCTCA
GCTTCCTCAGCCGCCGCCGCAGGCACAGCCGCTGCTGCCTCAGCCGCAGCCGCCCCCGCCGCCGCCCCCGCCGCCACCCGGCCC
GGCTGTGGCTGAGGAGCCGCTGCACCGACCgtgagtttgggcccgctgcagctccctgtcccggcgggtcccaggctacggcggggatggcggtaaccc
tgcagcctgcgggccggcgacacgaaccccggccccgcagagacagagtgacccagcaacccagagcccatgag**ggaca**cccgcccccctcctggggcgaggcctt
cccccacttcagccccgctccctcacttgggtcttcccttgtcctctcgcgaggggaggcag**agccttgttggggcctgtcctgaattcgat**ggccctcgtcttgcggggtc
tctggcctccctcagagg**agaca**gagccgggtcaggccagccagggactcgctgaggggcgtcacg**actccagtgccttcgccgttcccagtttgc**gaagttagggaa
cgaacttgtttctctcttctggagaaactggggcggtggcgcacatgactgttgtgaagagaacttggagaggcagagatctctagggttacctcctcatcaggcctaag
agctgggagtgca**ggaca**gcgtgagagatgtgcgggtagtggatgacataatgcttttaggaggtctcggcgggagtgctgagggcgggggagtgtgaacgcatcca
atgggatattctttttccaagtgacacttgaagcagcctgtgactcgaggcacttcgtactctcctggcgtttcatttagtttgtggtgtagtg**tagttaaaccaggttttaa**
**gcatagccaga**gaggtgtgcttctgtgtgtctgcaggcagttggatgagttgtatttgtcaagtacatggtgagttacttaggtgtgattattaataaaaaactatatgtgt
gcatatatatgaaagagtcgacttatacttaactgcctatcgatttttttgttctatataaaacggatacattggtggtgctcagttttcaccggggaatgaattttactagtg
ttgcagacaggcttgttttagaacataggccactctgactctgactttgtgccagtaaaagttcctgtttagttctttgctgacatcttatagatctttggaagctagctgctt
gtgactggagagaatattgaaacagaagagagaccatgagtcacagtgctctaagagaaaagacgctcaaaacatttcctggaaatccatgctgagtgttgagccctg
tgctctcttgcagctcagtcctttctctcaactctgggcattttatttctaatctggatttgtataattaataaggagaacttttgggaacaacctactaaagaatgtcatcat
taaaactcacttagaaaataagt

-Human insert sequence
-Mouse sequence
- m6A motifs
- m6A ACA sites analyzed by MazF-qPCR
- cryptic polyA1 site according Sathasivam *et al* [5]
- cryptic polyA2 site according Neueder *et al* [4]
- gRNA3
- gRNA2
- gRNA1

**B** **Targeted m6A sites analzyed by MazF-qPCR of the gRNAs used in CRISPR approach**

| | | |
|---|---|---|
| gRNA1 | Upstream | GGACA hm: 560 nt<br>GGACAhum: 399 nt<br>GGACAms: 186 nt |
|  | Downstream | ----- |
| gRNA2 | Upstream | GGACAhum: 207 nt<br>GGACAms: 46 nt |
| | Downstream | GGACAms: 167 nt |
| gRNA3 | Upstream | GGACAhm: 90 nt |
| | Downstream | AGACAms: 70 nt<br>GGACAms: 284 nt |

**Supplementary Figure 4. *Htt* RNA sequence of chimeric ST*Hdh*$^{Q111/Q111}$ cells/ *Hdh*$^{+/Q111}$ mice and potential DRACH motifs targeted by the CRISPRdCas13 approach. (A)** FASTA sequence of mutant *Htt* RNA (5' → 3') of ST*Hdh*$^{Q111/Q111}$ cells/ *Hdh*$^{+/Q111}$ mice. Sequence of the human insert is shown in orange and murine sequence in black. The indicated colors in the figure highlight m6A motifs in the first 524bp of intron1; the m6A-ACA sites analyzed by MazF qPCR; cryptic sites and gRNAs used for targeting. **(B)** Table showing the location of the target m6A sites analyzed by MazF qPCR for each gRNA used in the CRISPR approach.

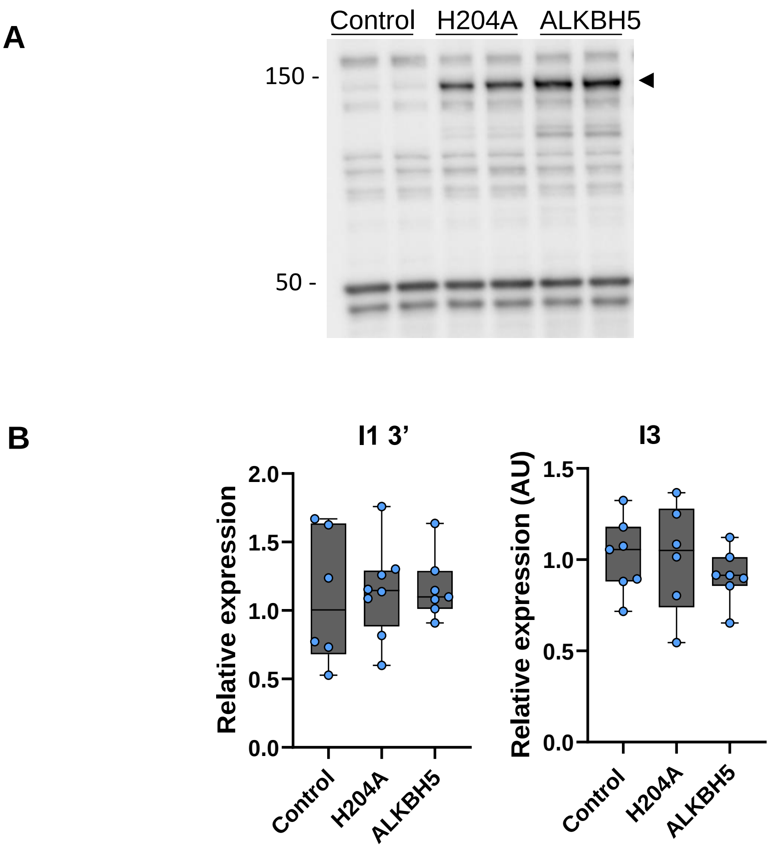


**Supplementary Figure 5. Fusion protein dCas13b-ALKBH5 is expressed in stable transfected ST*Hdh*$^{Q111/Q111}$ cells and only affect *Htt1a* expression (A)** Representative Western Blots showing expression of fusion protein dCas13b-ALKBH5 (A5) in cells stably transfected with dCas13b-A5 NT-gRNA (control), dCas13b-H204 gRNA2 and dCas13b-A5 gRNA2. **(B)** Expression levels of I1-3' and I3 *Htt* transcripts in transfected ST*Hdh*$^{Q111/Q111}$ cells with dCas13b-NT gRNA, dCas13b-H204 gRNA2 and dCas13b-A5 gRNA2. (n=4 independent experiments)

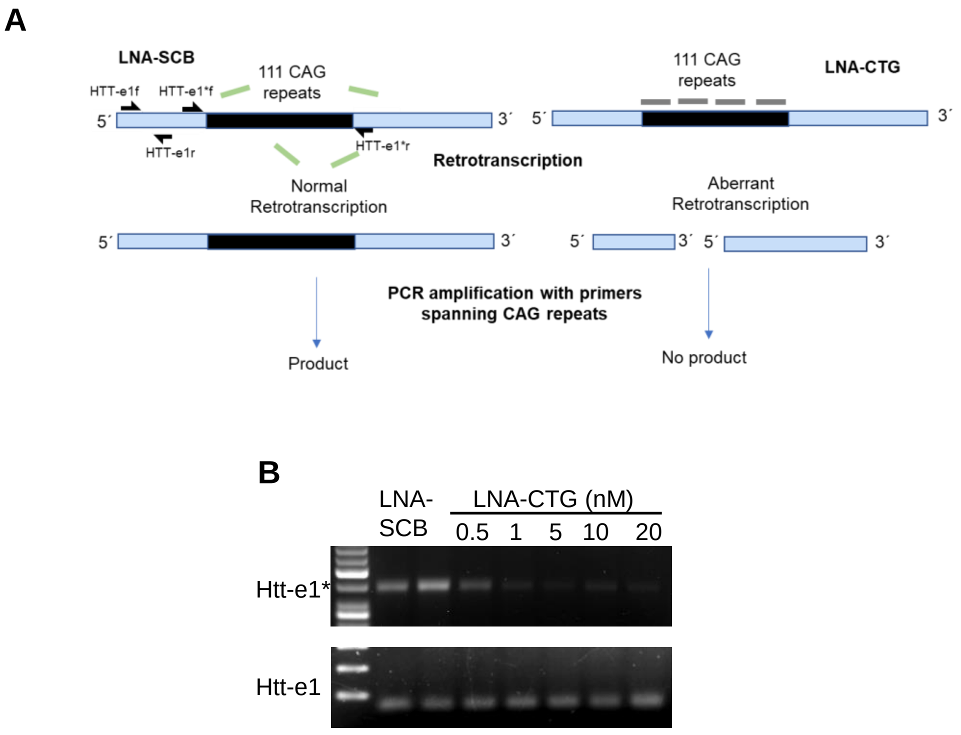


**Supplementary Figure 6. LNA-CTGs bind to *HTT*-e CAG repeats (A)** Scheme showing LNA-CTGs binding determination by the lack of PCR amplification within the LNA-bound region due to the strong incompatibility of LNA-CTG:CAG duplexes with retrotranscription and subsequent PCR amplification. *HTT*-e1binding sites of the primers used for PCR amplification in *HTT* exon 1 (*HTT_e1** and *HTT_e1* sets of primers) are shown **(B)** Gel electrophoresis showing *HTT* RT-PCR products from ST*Hdh*$^{Q111/Q111}$ cells transfected with different concentrations of LNA-CTG or LNA-SCB using primer sets HTT-e1*(for amplification of CAG expansions) and HTT-e1 (for amplification at 5´ of CAGs expansions)
